# Personalized and graph genomes reveal missing signal in epigenomic data

**DOI:** 10.1101/457101

**Authors:** Cristian Groza, Tony Kwan, Nicole Soranzo, Tomi Pastinen, Guillaume Bourque

## Abstract

**Background:** Epigenomic studies that use next generation sequencing experiments typically rely on the alignment of reads to a reference sequence. However, because of genetic diversity and the diploid nature of the human genome, we hypothesized that using a generic reference could lead to incorrectly mapped reads and bias downstream results.

**Results:** We show that accounting for genetic variation using a modified reference genome (MPG) or a *denovo* assembled genome (DPG) can alter histone H3K4me1 and H3K27ac ChIP-seq peak calls by either creating new personal peaks or by the loss of reference peaks. MPGs are found to alter approximately 1% of peak calls while DPGs alter up to 5% of peaks. We also show statistically significant differences in the amount of reads observed in regions associated with the new, altered and unchanged peaks. We report that short insertions and deletions (indels), followed by single nucleotide variants (SNVs), have the highest probability of modifying peak calls. A counter-balancing factor is peak width, with wider calls being less likely to be altered. Next, because high-quality DPGs remain hard to obtain, we show that using a graph personalized genome (GPG), represents a reasonable compromise between MPGs and DPGs and alters about 2.5% of peak calls. Finally, we demonstrate that altered peaks have a genomic distribution typical of other peaks. For instance, for H3K4me1, 518 personal-only peaks were replicated using at least two of three approaches, 394 of which were inside or within 10Kb of a gene.

**Conclusions:** Analysing epigenomic datasets with personalized and graph genomes allows the recovery of new peaks enriched for indels and SNVs. These altered peaks are more likely to differ between individuals and, as such, could be relevant in the study of various human phenotypes.

## Background

Standard ChIP-seq analysis relies on aligning reads to a reference sequence followed by peak calling [1] [2]. While the reference genome is a good approximation of the sequence under study, it does not account for the millions of small genetic variants, the larger structural variants or the two haplotypes of the human genome [3]. Instead, aligners cope with variation by allowing mismatches and indels in read alignments [4]. For example, reads that align to the SNP shown in Fig 1a would simply include a mismatch in their alignment to the reference sequence. Differences between the genome under study and the reference will shift the mapping of certain reads and generate unmapped reads (Fig 1a), a phenomenon known as reference bias [5]. Provided that the mapping of a number of reads is modified, an alignment to a personalized genome could lead to the gain or the loss of a peak, or what we will call an altered peak (AP). Actually, it has already been shown that just changing the assembly version of the reference can affect epigenomic analyses [6].

**Figure 1:**
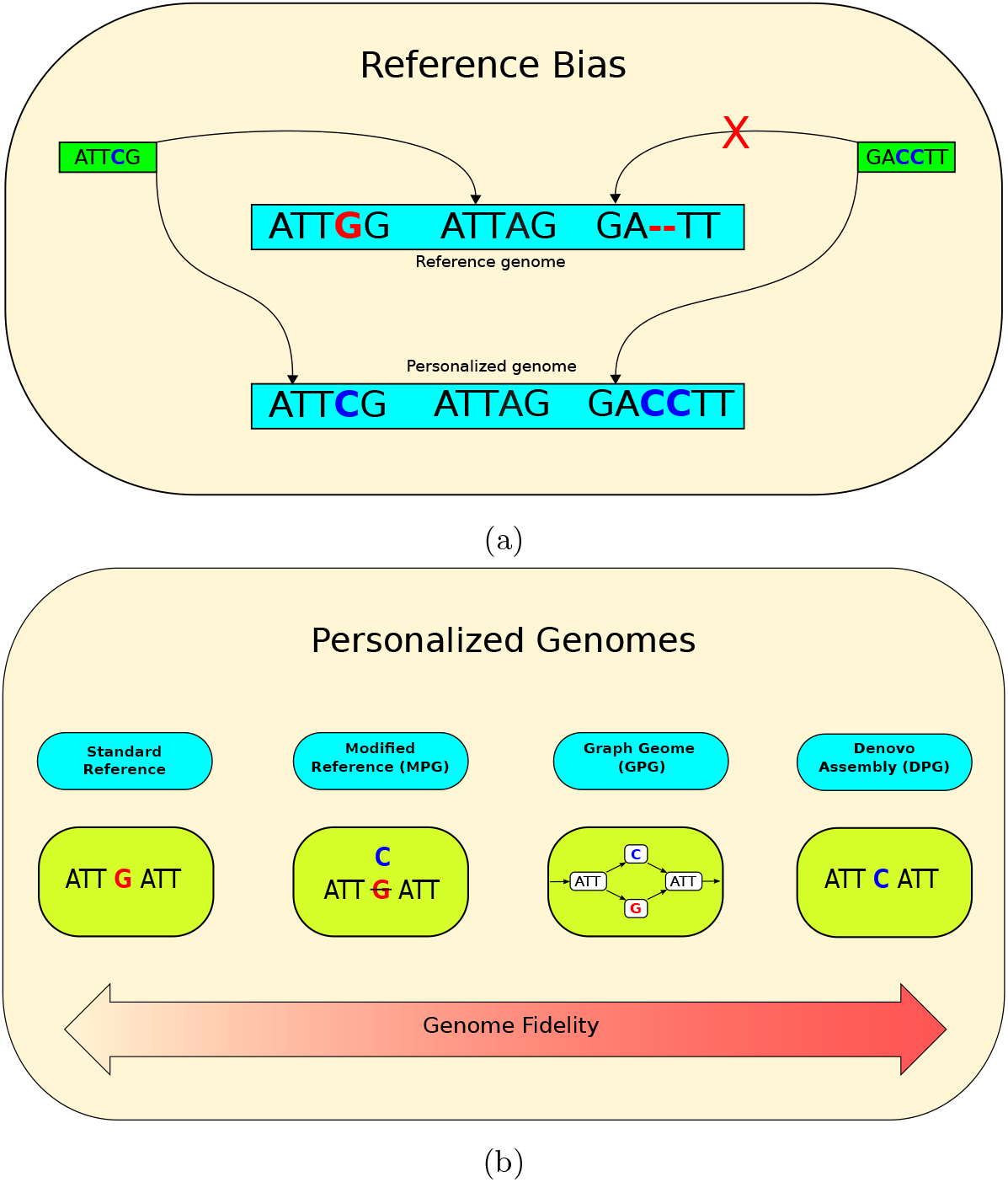
a) Two instances of reference bias that could be corrected by a personalized genome. One read is mapped to the incorrect location in the reference genome. The other read is unmapped in the reference genome, but becomes mapped in the personalized genome. b) Personalized genomes can be implemented in several ways. The reference can be patched with called variants to create a modified personal genome (MPG). Alternatively, a sequence graph genome could be augmented with an individual’s alleles (GPG). Finally, the entire personal genomic sequence can be assembled denovo (DPG).

In the current study, we wanted to evaluate the impact of using different types of personalized genomes on ChIP-seq analysis (Fig 1b). One obvious way of generating a personalized genome is to modify the reference genome using phased variant calls obtained from whole-genome sequencing to generate a diploid pair of sequences [7]. We call this making a modified personalized genome (MPGs). Because we cannot align reads to both MPGs simultaneously, analyses are done separately for each haploid sequence and merged afterwards. The advantage is that aligned reads would no longer feature the mismatch corresponding to the SNP mentioned above (Fig 1b). Epigenomic studies involving the use of MPGs are present in the literature. For instance, Shi et al. modified the reference genome using phased single nucleotide variant (SNV) calls and then realigned transcription factor and histone ChIP-seq data to record allelic specific binding events [8]. However, that study did not consider indels and was limited to understanding how SNVs affect standard analyses but not the identification of APs. Additionally, although pipelines such as AlleleSeq [7] do support indels and structural variations (SVs), they remain restricted to detecting allellic specific events without providing a way to detect APs. Allim [9] is a similar pipeline that attempts to detect instances of allelic imbalance in gene expression by modifying the reference to construct parental haplotypes. Turro et al. also leveraged genotypes, this time by modifying a reference transcriptome [10]. A study that did look at the use of MPGs as compared to the reference genome was done in the context of RNA-seq [11], where it was shown that personalized mouse genomes can improve transcript abundance estimates.

Improving the reference using SNVs and indels can help account for variation of small length, but not for larger SVs. For this reason, we also turn to *denovo* assembled personal genomes (DPGs) to fully reconstruct the genome sequence under study and to capture a broader range of genetic differences (Fig 1b). However, high-quality DPGs remain challenging to obtain for epigenomic analyses, as they typically require at least 50X sequencing depth and long-reads, which remain costly [12]. Also, the computational time for DPGs is much higher than aligning to a reference and calling variants [13]. Moreover, denovo assemblies may contain defects and are often incomplete compared to the reference [14]. Despite this, they may still provide a useful point of comparison.

Finally, the above trade-offs also motivate the exploration of graph personalized genomes (GPG) as an additional strategy (Fig 1b). GPGs can leverage available call sets that include a broad range of variants, from SNPs and indels to catalogues of sequence resolved SVs and also capture the diploid nature of the human genome [15]. Genome graphs abandon the flat structure of the standard reference genome. Instead, the contigs of the genome are chunked into nodes and connected by edges [16]. Different alleles are then represented as additional nodes, providing alternative paths for read alignment. Well-defined rules about the semantics of nodes and edges allow graphs to express all kinds of genetic variants in a concise manner, from small sequence differences to genomic rearrangements. Therefore, reads aligning to the SNP would take the path through the node that represents it. Conveniently, genome graph implementations such as `vg` [17] provide the proper utilities and semantics to work with annotations spanning multiple coordinate systems. Moreover, there are tools that can call ChIP-seq peaks directly from graph genomes [18].

The objective of our study is to provide a comparison between alternative personalized genomes (MPGs, DPGs and GPGs) for ChIP-seq analyses. Even if only a fraction of peaks are observed to be altered, these regions will correspond to biochemically active regions that are more likely to differ between individuals and, as such, could be relevant in the study of various human phenotypes.

## Results

### Modified personal genomes alter a small fraction of peaks that are enriched in indels

There are many high-confidence variant call sets and assemblies of the NA12878 genome, which makes it a good candidate for benchmarking [19] [20]. We created a paternal and maternal MPG for NA12878 and aligned whole-genome sequencing (WGS) reads to the standard human reference and to these MPGs (Methods). We wanted to estimate the proportion of changed mappings and noted that 3.6% of WGS reads move depending on the reference that is used (Table S1a). To measure the impact of reads changing location on ChIP-seq calls, we aligned H3K4me1 and H3K27ac ENCODE datasets from NA12878, and counted the proportion of altered peaks (Methods). We found that the fraction of personal-only and ref-only peaks was consistent between the two histone marks (Table 1). Among the H3K4me1 calls, each MPG yielded roughly 1600 personal-only (1.1%) peaks and roughly 800 ref-only peaks (0.6%). Among the H3K27ac calls, we called roughly 600 personal-only peaks (1.0%) and 300 ref-only peaks (0.5%) in each MPG. Notably, personal-only peaks were found at about double the rate of ref-only peaks. Ref-only peaks arise when the reads forming a peak pileup in the reference map to different locations in the personalized genome. In contrast, personal-only peaks emerge when reads shift their mapping from the reference pileup to the new personalized pileup or when reads that did not map to the reference become mapped to the personalized genome. Consistent with this hypothesis, there was a net gain of mapped WGS reads in the NA12878 MPG (Table S1a) and personal-only intervals are enriched in ChIP-seq rescued reads relative to ref-only intervals (Fig S1a).

**Table 1:** Number of altered peak calls in MPGs, DPGs and GPGs for the NA12878 H3K4me1 and H3K27ac marks.

| Version | Mark | common | personal-only | ref-only |
| --- | --- | --- | --- | --- |
| MPG, Paternal | H3K4me1 | 146520 | 1636 (1.1%) | 854 (0.6%) |
| MPG, Maternal | H3K4me1 | 146570 | 1622 (1.1%) | 808 (0.6%) |
| DPG, Hap1 | H3K4me1 | 141444 | 7176 (4.8%) | 6755 (4.6%) |
| DPG, Hap2 | H3K4me1 | 141442 | 7130 (4.8%) | 6774 (4.6%) |
| DPG, Pendleton | H3K4me1 | 142347 | 16245 (10.2%) | 8912 (5.8%) |
| GPG | H3K4me1 | 132668 | 3068 (2.3%) | 1178 (0.9%) |
| MPG, Paternal | H3K27ac | 68888 | 660 (1.0%) | 351 (0.5%) |
| MPG, Maternal | H3K27ac | 68909 | 688 (1.0%) | 335 (0.5%) |
| DPG, Hap1 | H3K27ac | 63419 | 2078 (3.2%) | 9901 (13.5%) |
| DPG, Hap2 | H3K27ac | 63441 | 2091 (3.2%) | 9899 (13.5%) |
| DPG, Pendleton | H3K27ac | 66811 | 5208 (7.2%) | 4980 (6.9%) |
| GPG | H3K27ac | 75538 | 1847 (2.4%) | 1206 (1.6%) |

Aligning to a personalized genome may cause differences in read density that do not necessarily lead to an AP call, especially in the strong peak regions. For that reason, we also counted the reads in personal-only, ref-only and common peak intervals and compared them between the reference and personalized alignments (Methods). Ideally, peaks that have AP calls should also have a skewed coverage. However, for most AP calls, we found that their coverage distribution remained clustered within the distribution of the common and unaffected peaks calls (Fig 2a). Most affected peak calls fall into the no-skew category together with common calls, with only around 30 peaks having a coverage skewed toward the reference or the MPG (Fig 2b and Table S2). Comparing the *q*-value distribution of common peaks against the distribution of APs revealed similar modes but a much shorter right tail for APs (Fig 2c). This means that personal-only peaks and ref-only peaks are confined to a region of narrower width and lower confidence than most common peaks (Fig S1b - S1c). Similar results were also observed for H3K27ac (Fig S2a and S3a).

**Figure 2:**
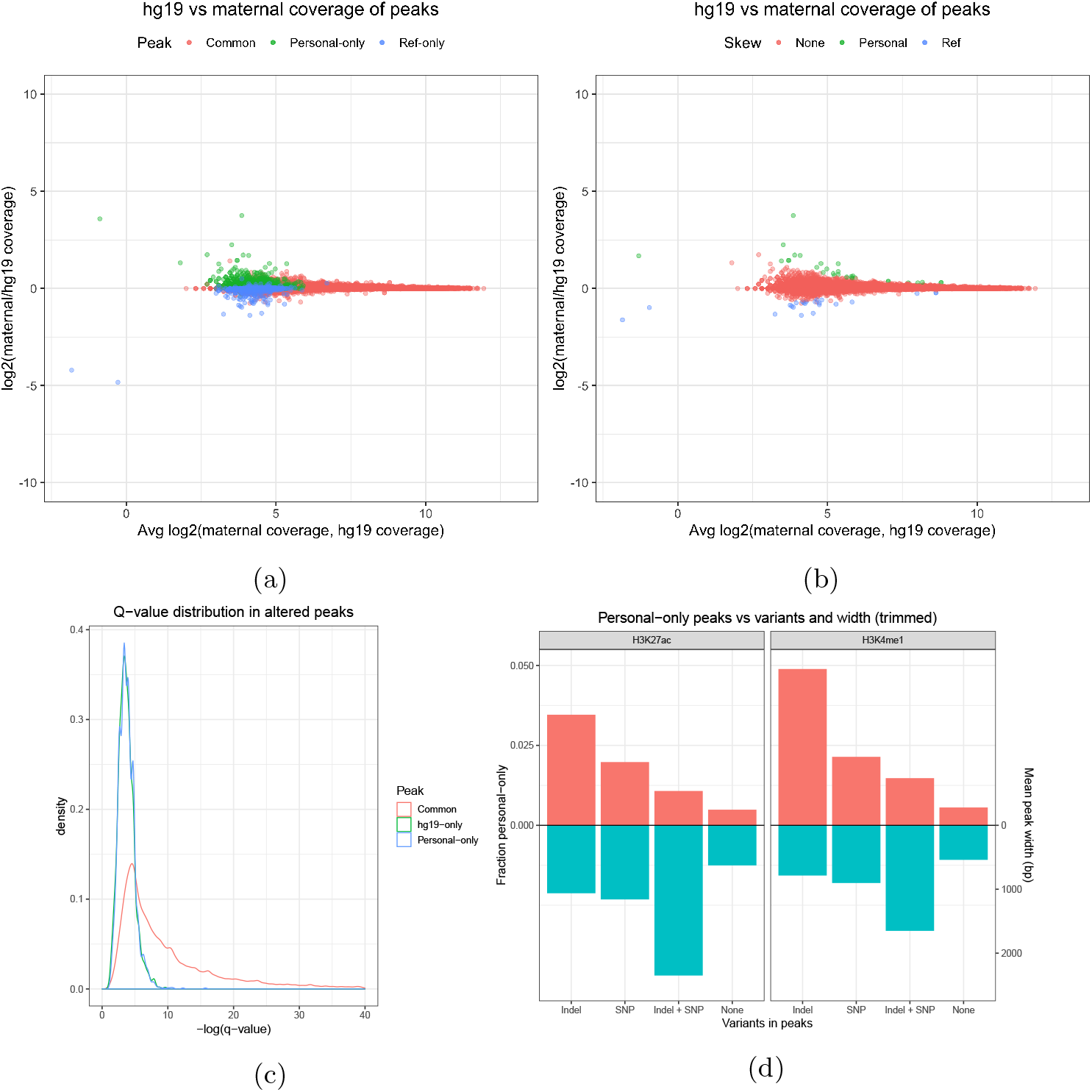
a) A comparison of the coverage of H3K4me1 peak called regions in hg19 and the maternal MPG. b) Identification of peak called regions that have a significant difference in coverage. c) Q-value distributions of the same H3K4me1 peaks. d) NA12878 MPG estimate of the probability that each combination of variation calls present in a region may cause a personal-only peak call compared to their average widths.

Finally, we wanted to explore the link between AP and variant calls, as we expected the former to occur mainly in the presence of the latter. For this purpose, we binned AP calls according to the overlapped combination of variants (Methods). Reassuringly, we found that peak calls that do not contain variations have a near zero chance of being altered, while peaks overlapping at least one indel are the most likely to be altered followed by peaks overlapping at least one SNP (Fig 2d). Interestingly, peaks containing at least one SNP and indel are the least likely to be altered. A factor that could explain this trend is the peak width associated to each peak category and histone mark. Indeed, we found that the mean width of peaks overlapping both indels and SNPs is the highest among the four combinations of variations, followed by peaks with at least one indel and peaks with at least one SNP (Fig 2d). Using a regularized logistic regression model (Methods), we were also able to show that peak width has an inverse relationship with AP calls (Fig S1d - S1e). We estimated that the AP call log-odds ratio decreases by 0.19 per additional 100 basepairs in peak width, increases by 1.29 per additional SNP and by 2.0 per additional indel.

### Applying modified personal genomes to Blueprint samples

NA12878 is a deeply sequenced sample with high quality variant calls, meaning that it is not representative of most datasets. We wanted to evaluate the proportion of altered peaks on lower pass WGS datasets such as Blueprint, a cohort of samples used in the study of haematopoietic epigenomes for which ChIP-seq data is available [21] (Methods). In Blueprint samples, we called on average 130 and 47 thousand common peaks for H3K4me1 and H3K27ac, respectively. Overall, the total number of peaks is comparable to NA12878 (Table S3). In H3K4me1, there are approximately 750 (0.6%) personal-only peaks and 450 (0.4%) ref-only peaks. In H3K27ac, there are approximately 330 (0.7%) personal-only peaks and 190 (0.4%) ref-only peaks. Among these samples, the number of APs is almost always below the NA12878 benchmark (Fig 3a and S4a). Again, ref-only peaks are observed to occur less often than personal-only peaks. A decrease is also observed with skewed peaks. While not numerous in the benchmark to begin with (50 to 70), their number in the typical Blueprint sample barely reaches double digits numbers (Fig 3b and S4b). This is likely due to the difference in the whole genome sequencing depth, as the NA12878 variant call set (3.5M SNPs, 0.5M indels) is richer than Blueprint (approximately 3.25M SNPs and 375M indels per sample, Fig S4c). We confirmed this by creating a NA12878 MPG by downsampling the original set to 2.6M SNVs and 100K indels. As shown in Table 1, the downsampled MPG produces fewer AP calls relative to the full set for both H3K4me1 and H3K27ac marks. We should also keep in mind that the phasing of NA12878 variant calls is better than for Blueprint, which could also contribute to more AP calls.

**Figure 3:**
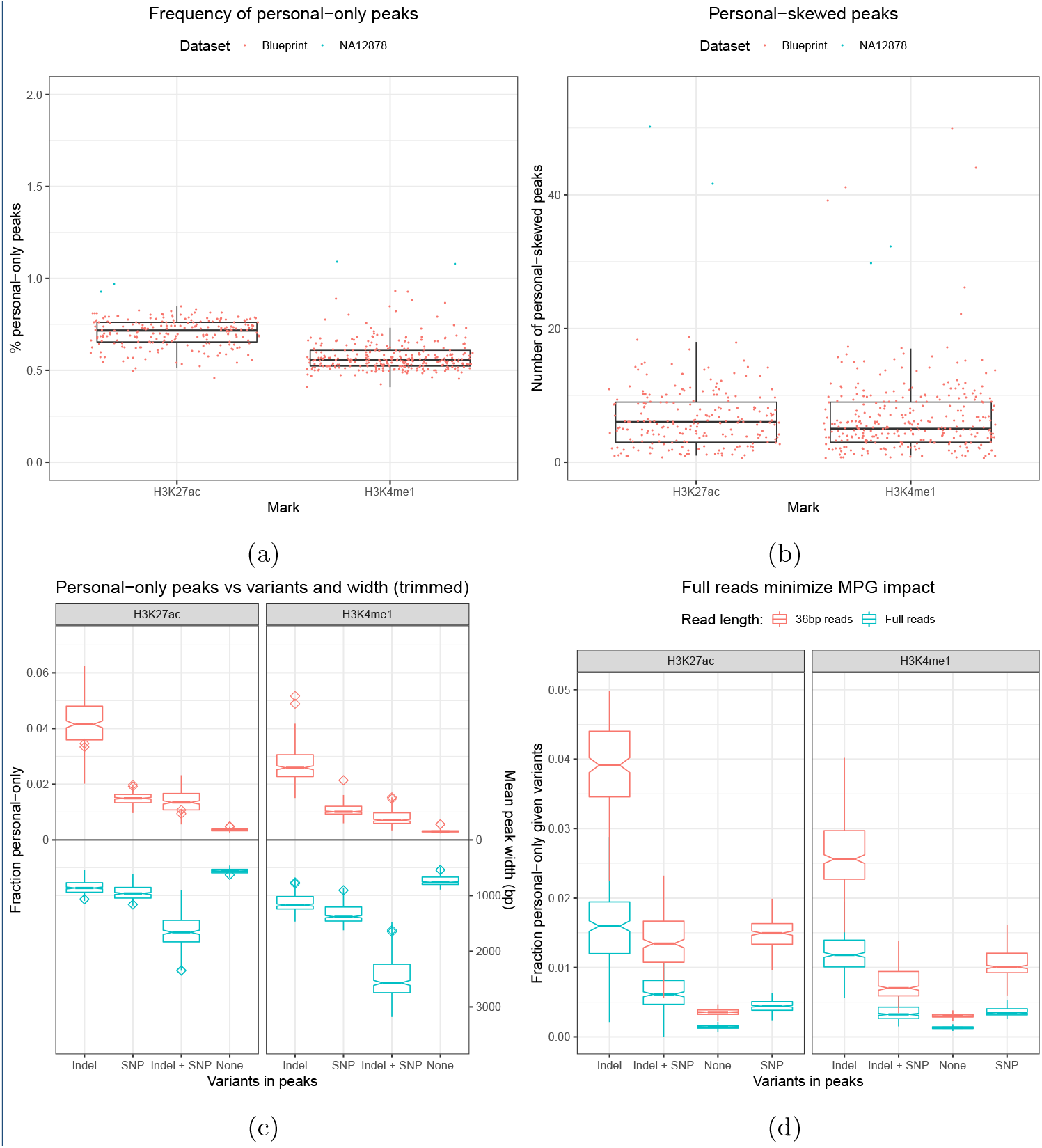
a) Proportion of peaks that are called only in personalized MPGs. b) Number of peaks with higher coverage in the personalized MPG than in the reference. c) Blueprint MPGs estimates of the probability that each combination of variation calls present in a region may cause a personal-only peak call compared to their relative average widths. d) The probability that a variant affects a peak called on full reads is lower compared to trimmed reads.

In Blueprint, altered peaks remain enriched in variants, with peaks containing indels being being altered most frequently (Fig 3c). Again, we found that the peaks of H3K4me1 were slightly less likely to be altered than the peaks of H3K27ac. As previously discussed, this is probably due to the inverse relationship between peak width and altered calls. As to the quality of altered Blueprint peaks, the same pattern of width, confidence and coverage observed in NA12878 were seen again in Blueprint samples (Fig S5). The small differences in coverage together with the weak confidence of APs, indicates that MPGs can only alter the calls of regions that are very near the threshold of significance.

Finally, in this initial analysis, we had trimmed every sample to a read length of 36 bp to make it comparable to the NA12878 datasets (Methods). To test the effect of read length, we repeated the Blueprint analysis with the full 100 bp reads. We found, as expected, that as the read length increases, APs become less likely (Fig 3d). We repeated the NA12878 WGS alignment comparison with the longer 100bp reads to gain some insight on why this happens (Table S4a). Compared to the shorter reads (Table S1a), the longer reads show a proportionally small decrease in aligned reads with unequal mappings. However, the proportion of reads that are mapped in one genome but not in the other halves. This is accompanied by a greater mapping rate of the whole WGS dataset. Therefore, the decrease in APs can be attributed to a smaller proportion of reads that are rescued by the personalized genome.

### Denovo personalized genomes create a larger number of altered peaks

If the moderate effect of using MPGs for ChIP-seq calls in NA12878 and Blueprint is explained by the fact that larger scale variations had not been taken into account, then denovo assemblies, or DPGs, could potentially have a broader impact. Support for this hypothesis comes from the increased rate of read mapping changes when using DPGs instead of MPGs (Table S1b). We opted to use the 10X Hap1 denovo assembly as a DPG for this comparison (Methods). In this DPG, 9.8% of reads change their mapping, which is nearly a three fold increase from the equivalent analysis with MPGs. When using full reads, we still get that 9.4% of reads alter their mapping (Table S4b). As in MPGs, the number of rescued reads proportionally changes the most.

In the context of ChIP-seq analysis, this should lead to a larger number of altered peaks. Indeed, using the same datasets (Methods), we found that the altered peak calls are roughly five times more numerous with a similar number of common peaks when using the Hap1 and Hap2 DPGs instead of an MPG (Table 1). For H3K4me1, we obtained approximately 7.1 thousand (4.8%) personal-only peaks and 6.7 thousand (4.6%) ref-only peaks. For H3K27ac, we called approximately 2.1 thousand (3.2%) personal-only peaks. For this mark, the number of ref-only peaks is unusually large at 9.9 thousand (13.5%) peaks. We also repeated the analysis that identifies peaks that have skewed read counts toward the DPG or the reference. Notably, we found that many AP calls now have substantial differences in coverage (Fig 4a and Fig S2c for H3K27ac). There are also many significantly skewed peaks, which are more distinguished in terms of reference and personalized read counts compared to their equivalents in MPGs (Fig 4b and Fig S2d for H3K27ac). Similar results are also obtained using the Pendleton DPG (Methods and Table 1). Overall, personal-skewed and ref-skewed peaks are one to two order of magnitude more numerous in DPGs versus MPGs (Table S2). Although personal-only peaks do not reach an identical distribution to common peaks, there are considerable gains in terms of width and quality (Fig S6a). DPG-only peaks are found to have a higher mean SNP and indel density compared to common peaks (Fig S6d). As for ref-only peaks, they are only slightly enriched in variation calls. This can be explained by a group of ref-only calls that have no coverage in the DPG and are probably completely missing from the denovo assembly (Fig 4a).

**Figure 4:**
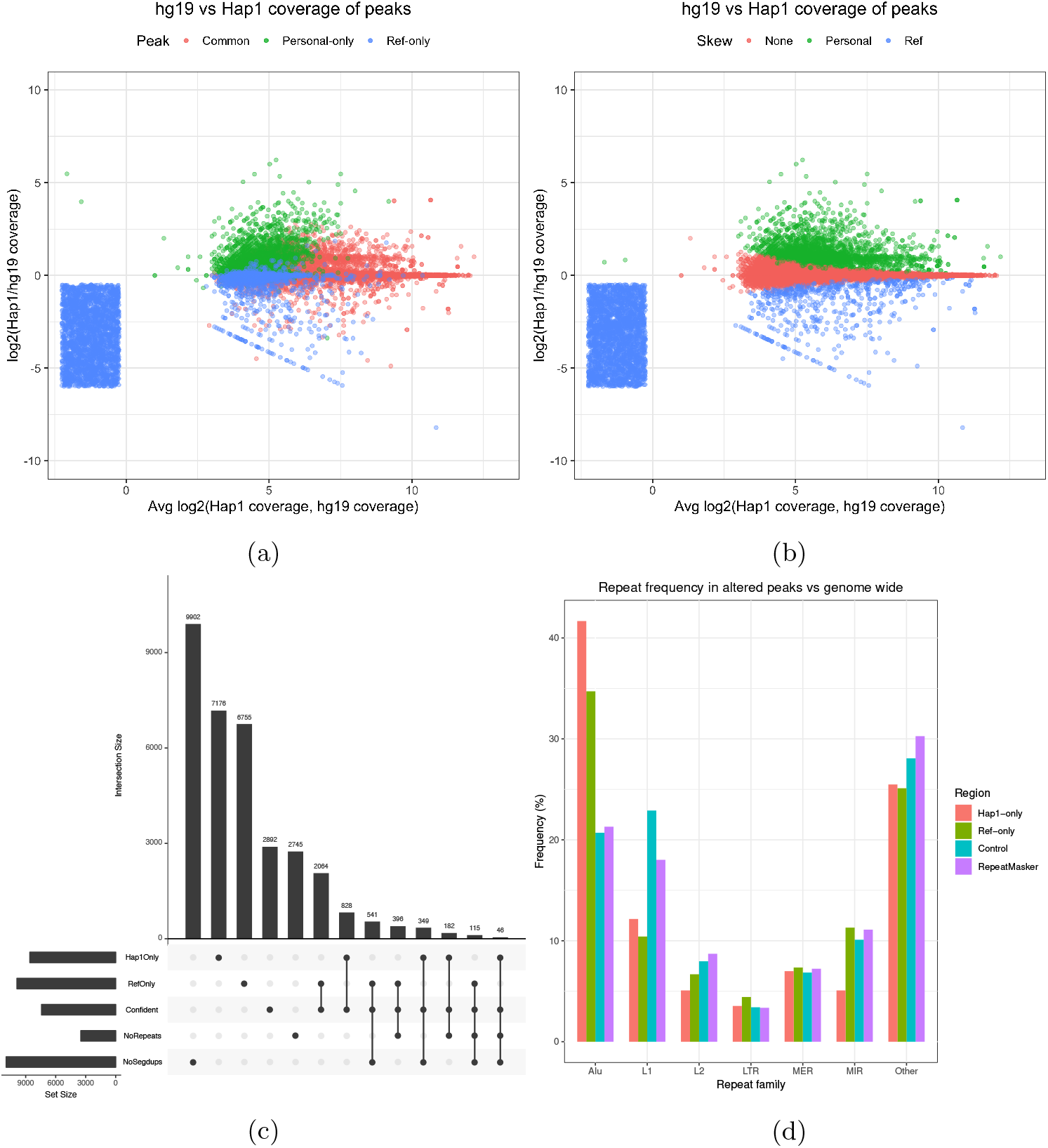
a) A comparison of the coverage of peak called regions in the reference and the Hap1 DPG. The smear represents ref-only peaks with no coverage in Hap1. b) Identification of peak called regions that have a significant difference in coverage. c) Summary of the overlap between altered peaks, confident peaks, repeats and segmental duplications [46]. d) The repeats that overlap altered peaks are enriched in Alu elements relative to their frequency in the RepeatMasker. The categories are chosen by grouping repeats by name prefix, summing their frequencies per group and taking the largest groups. Remaining groups are labeled as “other”. The control regions are random genomic intervals with a width distribution identical to altered peaks.

If SVs are the root of many AP calls, then many of these peaks should overlap repeats or segmental duplications that are known to be underrepresented in denovo assemblies [22]. We selected the most confident subset of H3K4me1 AP calls to be overlapped with SD and repeats annotations (Methods). This reduces the initial set to 828 confident DPG-only and 2064 confident ref-only peaks. Among confident DPG-only peaks, only 349 peaks are located in regions free of SDs (Fig 4c). Ref-only peaks with positive DPG coverage register much fewer SDs (6.3%). However, ref-only peaks without DPG coverage are highly associated with SDs (71.3%) (Table S5). The lack of coverage suggests that these duplicated sequences are not present in the DPG. Looking among the SD free peaks, we discovered peaks with large differences between the reference alignment and the DPG alignment (Fig S7). We also measured the enrichment in APs of the different repeat families (Methods). Alus were found to be 2 times more frequent in DPG-only peaks and 1.5 times in ref-only peaks (Fig 4d). The same is not true for repeat families such as L1, which occur equally or less often in APs relative to the genome. There also exists a small confident subset of 46 DPG-only and 115 ref-only peaks that are free of both SDs and repeats. Despite the absence of known repeats or segmental duplications, these peaks can still have large differences in coverage between the DPG and the reference alignments (Fig S8). We obtained similar results for H3K27ac (Fig S9a - S9b).

### Graph personalized genomes create more altered peaks than MPGs

Although DPGs are more effective than MPGs to recover APs, in practice they are often difficult to obtain. Therefore, we were interested in GPGs due to their ability to represent genetic variation and potentially approximate denovo assemblies by exploiting structural variant catalogues. In addition, GPGs improve on MPGs by allowing read alignment to a diploid genome instead of treating each haploid individually. As before, we mapped the same WGS reads to the reference genome, this time represented as a graph, and to the NA12878 GPG and then compared their coordinates using built-in `vg` functionality (Methods). By properly representing the diploid genome, we expected GPGs to shift the mapping of a greater proportion of reads than an equivalent pair of MPGs. In fact, we found that the proportion of unequal mappings between the reference graph and the NA12878 GPG (8.3%) is more than twice the number between the reference and the NA12878 MPGs (3.43%) given the same WGS dataset (Table S1c).

We found similar numbers of common peaks in GPGs as in MPGs and DPGs, specifically 132 thousand H3K4me1 calls and 75 thousand H3K27ac calls (Table 1 and Methods). Among the H3K4me1 calls, 3068 (2.3%) are personal-only and 1178 (0.9%) are ref-only. Among the H3K27ac calls, 1847 (2.4%) are personal-only and 1206 (1.6%) are ref-only. Both sets of values are intermediate between MPGs and DPGs (Table 1). Revisiting the peak read counts between the reference graph and the diploid graph shows greater dispersion, among both altered and common peaks (Fig 5a). The same test for read count skew yields an order of magnitude more peaks (279 to 411) than MPGs (Fig 5b, Table S2). See also Fig S2e - S2f for similar results with H3K27ac. Next, we recalculated the association of indels and SNPs with the personal-only peak calls in GPGs (Fig S10b). Again, indels have the strongest association with APs for both H3K4me1 and H3K27ac marks. Contrary to MPGs, H3K27ac peaks containing both indels and SNPs are just as likely to be altered as peaks containing only SNPs, despite being much wider. Similarly to MPGs, peaks lacking variants are the least likely to be altered in both histone markers.

**Figure 5:**
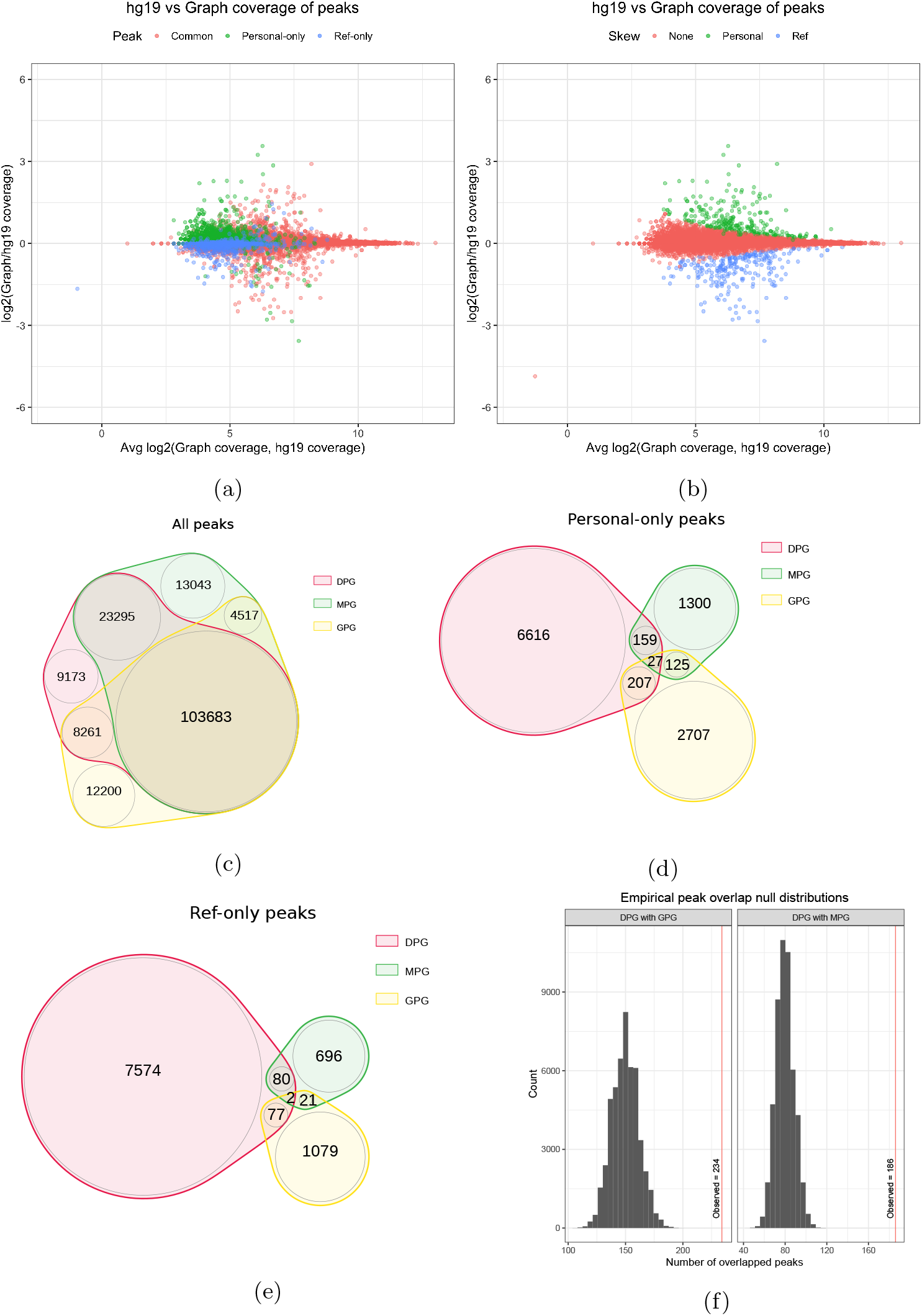
a) A comparison of the coverage of H3K4me1 peak called regions in the reference and the graph genome. Pairwise overlaps between MPG, DPG and GPG H3K4me1 peak tracks. b) Identification of peak called regions that have a significant difference in coverage. c) Overlap of all peak calls. d) Overlap of altered personal-only peak calls. e) Overlap of ref-only peak calls. f) Empirical null distributions for the overlap of personal-only peaks between personal genome implementations.

Next, we were interested between in the concordance between the 3 approaches: MGP, GPG and DPG (Methods). We found the overlap between the total peak tracks to be substantial, with over 100 000 H3K4me1 peak calls overlapping between the three personalized genome implementations (Fig 5c and S9d for H3K27ac). In contrast, when the AP calls are intersected, a small overlap is observed for personal-only peaks and ref-only peaks (Fig 5d - 5e and S9e - S9f for H3K27ac). Only 234 of 3068 (7.6%) of the NA12878 GPG personal-only calls are replicated in the DPG. Similarly, only 79 GPG ref-only calls are replicated from a total of 1178 peaks (9.8%). Comparatively, the replication rates between MPGs and DPGs are slightly higher, despite smaller absolute number of peaks. 186 of 1622 (14%) personal-only peaks and 82 of 808 (10.1%) ref-only peaks are replicated in the DPG. We wanted to know if chance alone could explain this small overlap of AP calls. We did this by generating a distribution of peak overlaps by randomly and repeatedly sampling the respective number of personal-only peaks in each genome from its total number of peaks (Methods). The expected number of replicated personal-only peaks is 140 peaks between the GPG and DPG and 80 peaks between the MPG and DPG (Fig 5f). As such, albeit small, the number of replicated peaks cannot be explained by chance alone.

### Further characterizing the altered peaks

We were interested in comparing the quality of the APs found by the three different approaches. We achieved this by comparing the q-values by rank in each genome (Fig 6a). From this, we observed that the best DPG-only peaks surpass the best GPG-only and MPG-only peaks by a wide margin. The top GPG APs only surpass the top MPG APs by around one unit on the –log_10_(*q*) scale. But on a linear scale, this means that the most confident GPG APs are an order of magnitude more confident than the most confident MPG APs. See Fig S3c - S3d for H3K27ac.

**Figure 6:**
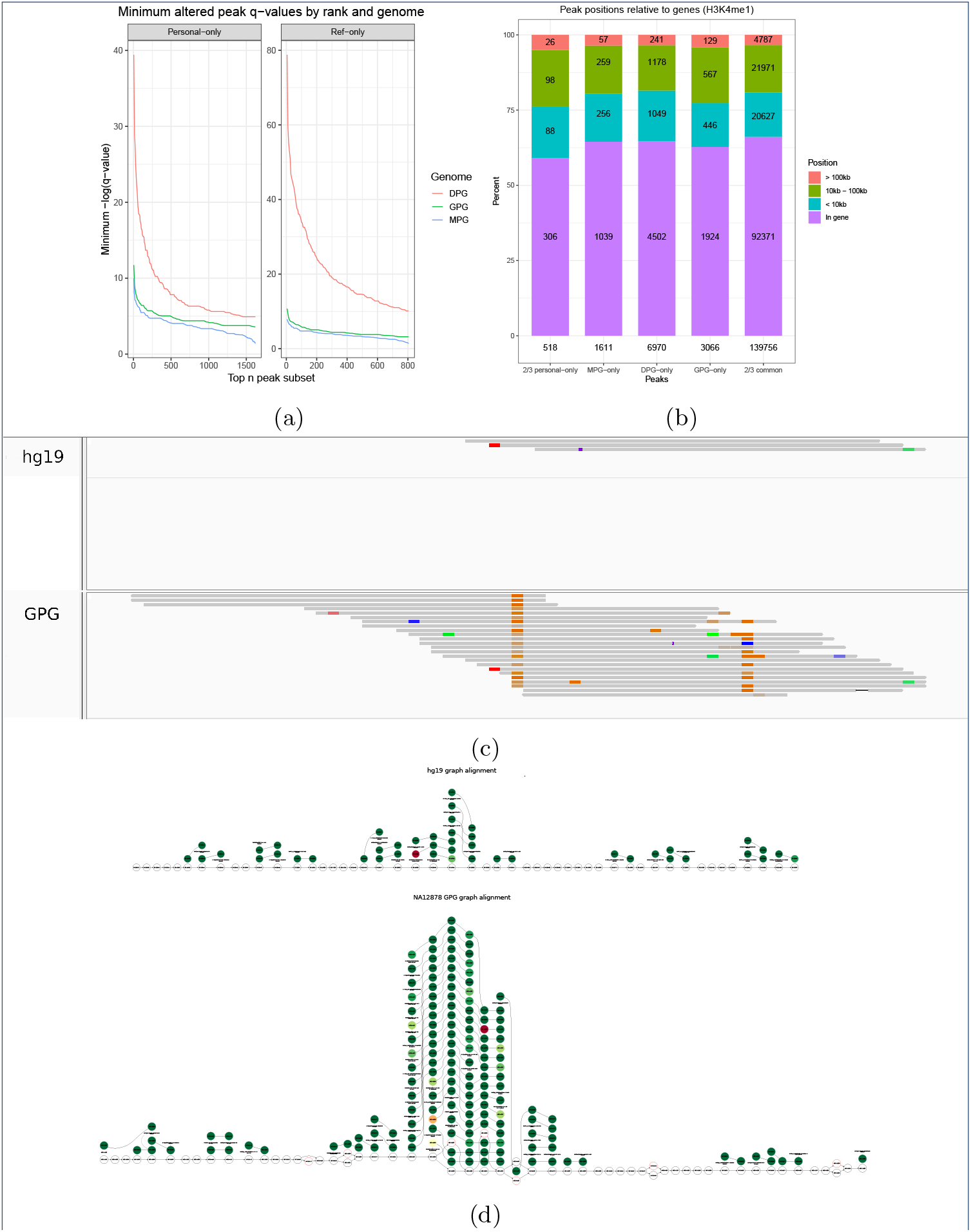
a) Comparison of altered peak q-values between MPG, GPG and DPG implementations by rank. The top n peak subset was increased by 5 peak increments. b) Distribution of gene relative positions of personal-only peaks among all genomes. Personal-only and common peaks replicated in at least two genomes are also featured. c) The pileup of a GPG-only peak projected to the hg19 linear reference. d) The true graph rendering of the above AP in the NA12878 GPG and reference genome graph.

Finally, H3K4me1 is a histone mark known to be associated with gene activation that is present near transcription start sites and transcribed regions [23]. However, this pattern may not necessarily be replicated in the AP calls, particularly if they are caused by noisy signal. Therefore, we wanted to check whether APs maintain the same genomic distribution as the rest of the calls, among all three genome implementations. To this end, we computed the distances to the nearest gene for MPG-only, DPG-only and GPG-only peaks as well as for personal-only and common peaks that were replicated in at least two genomes (Methods). We distinguished between peaks that overlap a gene and peaks that are within 10kb, between 10kb and 100kb, or further than 100kb from a gene. Overall, the genomic profile of AP calls is very similar to that of replicated common calls across the board, regardless of the genome or replication (Fig 6b and Fig S9c for H3K27ac).

Given that more than half of APs are within genes, some may be of particular interest. Indeed, Fig 6c shows a GPG-only example projected to the reference, while Fig 6d shows the true graph rendering of the pileups. The personalized peak overlaps four consecutive SNVs which are incorporated in the GPG but not the reference graph. The graph rendering clearly shows a fair number of reads aligning to these SNVs, forming a pileup that fails to appear in the reference graph. Moreover, this interval is within the third intron of STON1-GTF2A1L, a gene that appears in two GWAS studies linking it to neovascular age-related macular degeneration [24] and polycystic ovary syndrome [25]. Examples like this suggest that GPGs may improve our understanding of gene regulation in individual genomes.

## Discussion

By moving from the reference sequence to a MPG, GPG, and DPG, the genome representation became richer by incorporating SNVs and indels, variants in the form of a diploid graph, and also larger structural variants. When reanalyzing ChIP-seq datasets using these personalized genome implementations, we were able to identify hundreds to thousands of APs. While most APs detected using MPGs had only marginal changes in coverage, the GPGs and DPGs, yielded tens to thousands of peaks with significant read count differences relative to the reference. Notably, we observed that indels followed by SNVs were enriched in APs and that there was an inverse correlation with peak width. We also observed that Alus were overrepresented in APs, a transposable element known to be active in the human genome [26] and with many polymporphic instances in the population. Although it is tempting to think that some of these APs might be driven by these polymorphisms, it would require additional validation as it could also be caused by errors in the personalized genomes that were used for the analysis.

Although the vast majority of common peaks were identified consistently by the 3 methods, only a minority of APs were found by 2 or more methods. This limited overlap might be a consequence of the fact that the genome implementations are technically very different from each other. For instance, only DPGs at this stage took into account SVs but, at the same time, some regions of the personal genome might be missing for the current DPG. GPGs represent a promising compromise between MPG and DPGs as they also have the ability to natively account for the diploid nature of the human genome. A natural extension will be to try to incorporate SVs into GPGs to see how it can further improve their performance. Furthermore, as pangenome graphs are created to capture all known variations [16], it might be possible to further improve on the current performance even without the need for personalized graphs per say.

Finally, even though we primarily focused on APs, we also encountered peaks that differed significantly in read counts. These skewed common peaks produced by personalized genomes should not be ignored, particularly when performing differential expression analysis between control and treatment groups. Even if the number of skewed peaks is generally smaller than APs, they remain important because such studies typically identify a small number of diferentially expressed regions. Therefore, the application of personalized genomes could reveal new data points or correct false positives.

## Conclusions

Analysing epigenomic datasets with personalized and graph genomes allows the recovery of novel ChIP-seq peaks many of which fall within genic regions and could differ between individuals. Although we focused this study on ChIP-seq, it is likely that these results will extend to other epigenomic assays such as ATAC-seq and whole-genome bisulfite sequencing. As we move towards profiling the epigenome of large human cohorts to study various phenotypes, it is likely that using personalized and graph genomes will reveal important loci that would have been missed otherwise.

## Methods

### Data

We selected NA12878 as a benchmark dataset due to the availability of phased variation calls from high coverage whole genome sequencing (200X) [19] in addition to several denovo assemblies. FASTQs for H3K27ac, H3K4me1 marks and a control (input) were downloaded from the ENCODE project [27]. The accession numbers for these samples are ENCFF000ASM, ENCFF000ASU and ENCFF002ECP respectively. To generate additional supporting results for ChIP-seq, we used a low pass NA12878 WGS dataset from IGSR [28] (SRR622461).

Samples from the Blueprint project [29] were also selected due to the availability of phased variation calls from low pass whole genome sequencing (8X) together with ChIP-seq datasets for the H3K4me1 and H3K27ac histone marks. In total, 151 H3K4me1 samples and 111 H3K27ac samples were used in this analysis.

### Preparing personalized genomes

Three different approaches were used to generate personalized genomes. First, `vcf2diploid` [7] was used to substitute the alternative sequence of the phased variation calls into the hg19 reference to create a MPG. The output are two FASTA files for each contig, forming the conventionally named maternal and paternal haplotypes. The contig FASTAs were concatenated according to their haplotype, resulting in one maternal and one paternal FASTA. It is to be noted that `vcf2diploid` does not process unordered contigs. Therefore, unordered contigs were removed from hg19 to ensure the same set of contigs between the standard and substituted versions. Also, `vcf2diploid` generates two chain files that allow the lifting of annotation tracks with coordinates in hg19 to the corresponding personalized haplotype using `liftOver`. This is necessary since the incorporated indels shift the coordinates of the maternal/paternal haplotype relative to hg19.

The second approach, applied only to the NA12878 dataset, consisted of using denovo assembled genomes from the Pendleton [30] and two 10X Genomics assemblies [20] to create two DPGs. The 10X Genomics assembly includes two pseudo-haps named Hap1 and Hap2 that will be used as a denovo assembled diploid genome. In the case of the denovo assemblies, the chain files had to be produced from a BLAT [31] alignment between the denovo assembly and hg19 with the UCSC tool set [32]. This allowed the lifting of annotation tracks from the denovo assembly to the hg19 reference. The performance of DPG and MPG chain files was compared through the proportion peaks that failed to lift. Note that hg19 contains alternative contigs that represent some loci multiple times. In denovo assemblies, we expect loci to be represented only once. Therefore, the above analysis was performed on a hg19 version that was stripped of alternative contigs.

To allow alignment to personalized MPG or DPG FASTAs using `bwa mem` [33], an index was created using `bwa index`. A FASTA index was also created using `samtools faidx` [34] to compute the new chromosome sizes.

The third approach involved creating a reference graph genome by converting the linear hg19 reference to a graph format. A copy of this graph was then augmented with NA12878 variant calls, which yields the GPG. This was done with `vg construct` [17]. `xg` and `GCSA2` graph indices were created with `vg index` to allow mapping reads with `vg map`.

### Aligning, peak calling and annotating

To remove any effect of read length, all reads were trimmed to 36bp using `trimmomatic` [35]. The trimmed reads were aligned using `bwa mem` to hg19 and each personalized haplotype FASTAs. After marking duplicates with `picard` [36], peak calling was done on the corresponding BAM files using MACS2 [37] with `--nomodel` and the `--gsize` parameter set to 80% of the assembly length. In graph genomes, peaks were called with `Graph peak caller` [18], a graph MACS2 implementation, by using the same genome size and the same fragment length parameter that was estimated by MACS2.

For each alignment, a coverage annotation was produced with `bedtools bamtobed` [38]. The output was a BED file listing all the aligned reads and their coordinates. Graph alignments (GAM) were surjected to BAM using `vg surject` and underwent the same procedure.

### Lifting annotations

In the case of DPGs, coverage and peak annotations were lifted from the DPG to hg19 using the tool `liftOver` [39]. In the case of MPGs, the annotations were lifted from hg19 to the MPG. Therefore, this required the lifting of variant call annotations to the MPG in addition to the peak call and coverage annotations.

The variant call annotation was first converted from the VCF format to BED, separated by phase and type (SNP vs indel) and then lifted to the personalized haplotype. The outcome is a set of BED files listing the SNPs and indels separately for each respective haplotype.

`liftOver` was called with default arguments in BP samples, which require 95% sequence identity between lifted regions and target regions. This stringent `-minMatch` was not an issue since MPGs are almost identical to hg19 and virtually all peaks lift. In 10X and Pendleton samples, `-minMatch` was set 0.85 to reduce the number of unlifted peaks and reduce the number of false ref-only peaks. To evaluate lifting efficacy, the number of peaks that failed to lift was compiled for every sample. Once tracks are lifted to a common coordinate system, it becomes possible to overlap and compare the annotations from the personalized haplotype and the hg19 standard reference using `bedtools` [38].

Graph annotations are readily surjected onto hg19 using built-in functionality in `vg` and `Graph peak caller`.

### Overlapping annotations

The lifted or surjected peak call annotations were overlapped using `bedtools intersect` and `bedtools subtract` [38]. Peaks resulting from the intersect of the personalized and the hg19 peak tracks were categorized as **common**. Peaks resulting from subtracting the hg19 track from the personalized track were categorized as **personal-only**. Similarly, peaks resulting from subtracting the personalized track from the hg19 track were categorized as **ref-only**. The end result is a set of three BED files for each personalized genome containing the common peaks, the personal-only peaks and ref-only peaks.

The number of variation calls in each peak was calculated. The corresponding indel and SNP tracks were intersected with the track of each category of peaks using `bedtools intersect -c` to list the number of variations overlapping each common, personal-only and ref-only peak.

Furthermore, the peak tracks were overlapped with the coverage tracks of the personalized and hg19 versions of the alignment using `bedtools intersect -c`. The output is the original peak track with an additional field listing the number of reads in each peak. As a result, the number of reads in regions corresponding to the peaks is known in the reference alignment and the personalized alignment.

### Finding peaks with skewed coverage

To find peak called regions that have significant differences between their hg19 and personalized coverages, a statistical test was needed. This comparison is similar to differential expression in that read counts are compared between two conditions: the hg19 reference and the personalized assembly. For the purpose of differential expression, technical variation that occurs during the preparation of different libraries is known to be underestimated by Poisson based tests (overdispersion) [40]. However, unlike differential expression, our read counts are not compared between multiple sequencing experiments done under the two conditions. Instead, there is only one dataset that was aligned to two different assemblies, which implies that biological and technical variation is not present here in the same way. Therefore, we simply used a *χ*^2^ test with a significance value *α* of 0.05 to detect peaks with skewed coverage. We obtained an identical result with the edgeR package [41] by setting the dispersion parameter to 1 × 10^-3^ (near 0). Peaks with null coverage in one of the alignment versions were artificially assigned one read to allow applying the test. Peaks with insignificant differences were placed in the no-skew category. An overview of the above steps can be found in figures S11a and S11b.

Peaks that had significant differences with a higher coverage in hg19 than in the personalized haplotype were categorized as ref-skewed. Similarly, peaks that had a higher coverage in the personalized genome than in the reference were categorized as personal-skewed.

### Characterizing altered peak calls

To quantify the fraction of AP calls, the number of ref-only and personal-only peaks was counted and then divided by the total number of peaks to obtain their frequency relative to the total number of peaks in their sample. For each sample, the set of all peaks was divided into mutually exclusive categories according to the combination of overlapping variation calls (SNPs only, indels only, SNPs and indels, none). The same was repeated for ref-only and personal-only peaks. For any given variation category, the counts of ref-only and personal-only peaks were divided by the sample wide peak count of the given category to obtain the probability that the peak call could be affected by that specific combination of variations. At the same time, the mean peak widths were recorded.

For DPGs, we counted the number of hg19-relative variant calls overlapping common, ref-only and personal-only peaks. We did this to check whether ref-only peaks and personal-only peaks remained enriched in hg19-relative variation calls compared to common peaks, despite the fact that they originate from peak calls in a denovo assembly and not hg19 itself.

We also counted the overlaps of altered peaks in DPGs with SDs and repeats from the RepeatMasker annotation. Repeats were first grouped by family. Confident peaks were selected by removing any peak with a log(MACS2 score) < 4.0. This value was chosen because it excludes uncertain and uninteresting peak calls and most APs generated by MPGs.

Logistic regression was performed on NA12878 H3K4me1 peaks with AP/common as a binary response variable and peak width, SNP count and indel count as covariates using the `glmnet` [42] R package. Ref-only and personal-only peaks were coded as AP=1 and common peaks were coded as AP=0. Lastly, common peaks were downsampled to the number of AP calls to avoid unbalanced classes. Since the fitting algorithm is non-deterministic, we ran `cv.glmnet` 1000 times and reported the median coefficient values.

### Comparing WGS alignments between genomes

If peak track differences occur between two assemblies, they should be corroborated by differences in the mapping of a sufficient number of reads between their raw alignments. That is, the proportion of reads with different mappings between the reference and the personalized genome should be considerable. To show this, we used `Jvarkit cmpbamsandbuild` [43] to compare the DPG and MPG alignments of the low pass NA12878 whole genome dataset to hg19. The same comparison was done between the reference and the paternal NA12878 MPG. To compare the GPG and the reference graph alignments, `vg gamcompare` was used instead. For unequal mappings, we considered reads that are mapped more than 100bp apart, reads that are mapped in one build but not the other, and reads that fail to lift between assemblies. We add these proportions to obtain the final proportion of changed mappings. The IGSR WGS dataset was chosen instead of a ChIP-seq dataset because we expect a more uniform coverage of genomic regions.

### Finding replicated peaks among MPGs, DPGs and GPGs

To get the replicated calls between the DPG and the MPG approaches, the personalized tracks needed to be lifted to a common coordinate system in hg19. This is necessary because the MPG APs were computed in MPG coordinates, while the DPG and GPG APs were computed in hg19 coordinates. To do so, chain files were created through the previous BLAT method to lift the MPGs to hg19. Once the tracks of ref-only and personal-only peaks respective to the MPGs was lifted to hg19, `GenomicRanges` [44] was used to calculate the pairwise overlap of peak calls between the three approaches and identify peaks that are replicated with at least two of the three methods. This package was also used to characterize the position of peaks relative to genes in the UCSC genes annotation. A Venn diagram was produced for personal-only calls, ref-only calls and all peak calls using `nVenn` [45].

## Ethics approval and consent to participate

Not applicable.

## Consent for publication

Not applicable.

## Availability of data and materials

The NA12878 ChIP-seq datasets are available in the ENCODE repository under accessions ENCFF000ASM, ENCFF000ASU and ENCFF002ECP.

The NA12878 variant calls are available at ftp://ussd-ftp.illumina.com/2017-1.0/hg19/small_variants/NA12878/.

The WGS dataset is available in the IGSR repository under accession SRR622461.

The 10X Genomics denovo assemblies are available at https://support.10xgenomics.com/de-novo-assembly/datasets/1.0.0/NA12878.

The Pendleton assembly is available in the BioProject repository under accession PRJNA253696.

The Blueprint data is available from the European Genome-phenome Archive upon application to the BLUEPRINT Data Access Committee, which we applied for and received access to. More information at http://dcc.blueprint-epigenome.eu/#/md/dac_applications.

Annotations are available from the UCSC Table Browser at https://genome.ucsc.edu/cgi-bin/hgTables.

## Competing interests

The authors declare that they have no competing interests.

## Funding

This work was supported by grants from the Canadian Institutes of Health Research (EP1-120608, EP2-120609, and CEE-151618).

## Authors’ contributions

CG performed the majority of data analysis, interpretation of results, and writing of the manuscript, including production of all the figures. TK contributed to manuscript preparation. NS contributed to data collection and TP contributed to the study design. GB designed and led the study and contributed to data interpretation, and manuscript preparation.

## Acknowledgements

We would like to thank Bing Ge who was involved in the variant calling of the Blueprint samples. We would also like to acknowledge Calcul Quebec and Compute Canada, for access to resources to perform these analyses.

## Supporting figures

**Figure S1:**
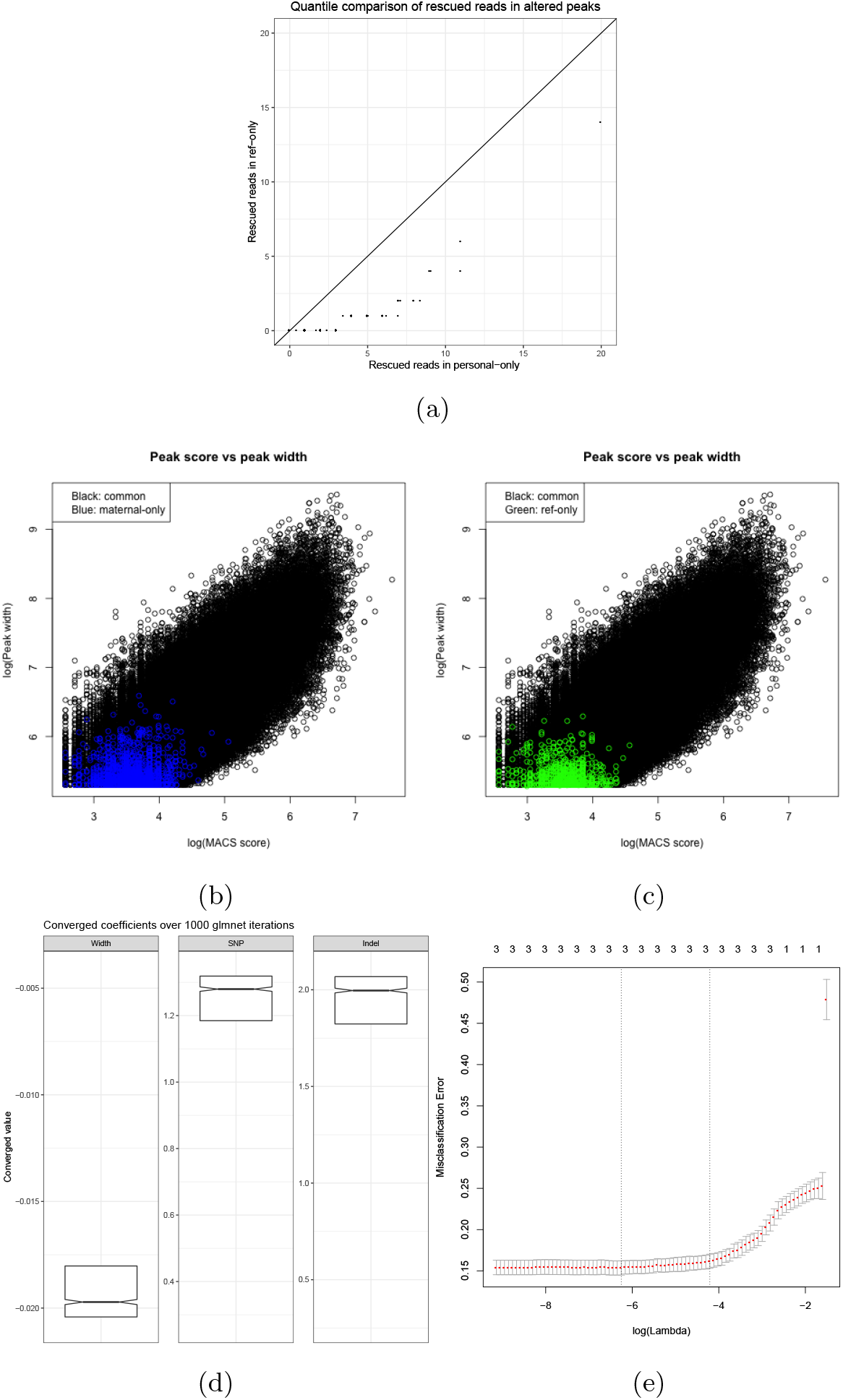
a) Quantile-quantile comparison between the distribution of rescued reads in ref-only and personal-only distributions. b) Confidence and width of the NA12878 maternal MPG altered peak calls (H3K4me1). Peaks called only in the maternal MPG against the common peak background. c) Peaks called only in hg19 against the common peak background. d) Distributions of converged coefficients for the width, SNP count and indel count terms in the `glmnet` logistic regression model. Median coefficients are 0.19 for width, 1.29 for SNP 1.9 for indels. e) The chosen regularization parameter corresponds to the minimum misclassificaton error during k-fold cross-validation. This model achieves a 0.15 misclassificaton error.

**Figure S2:**
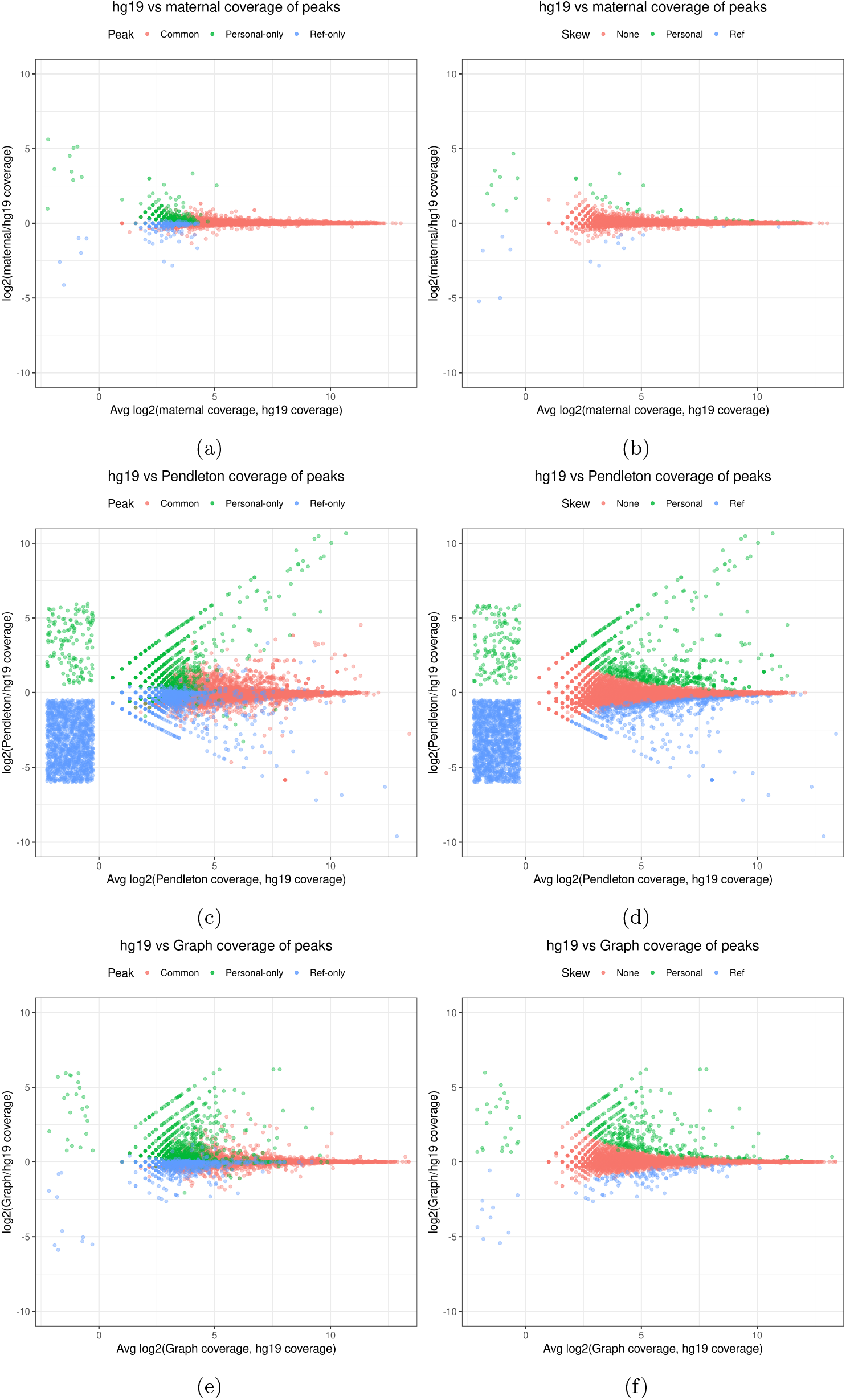
MA plots for H3K27ac analogous to H3K4me1.

**Figure S3:**
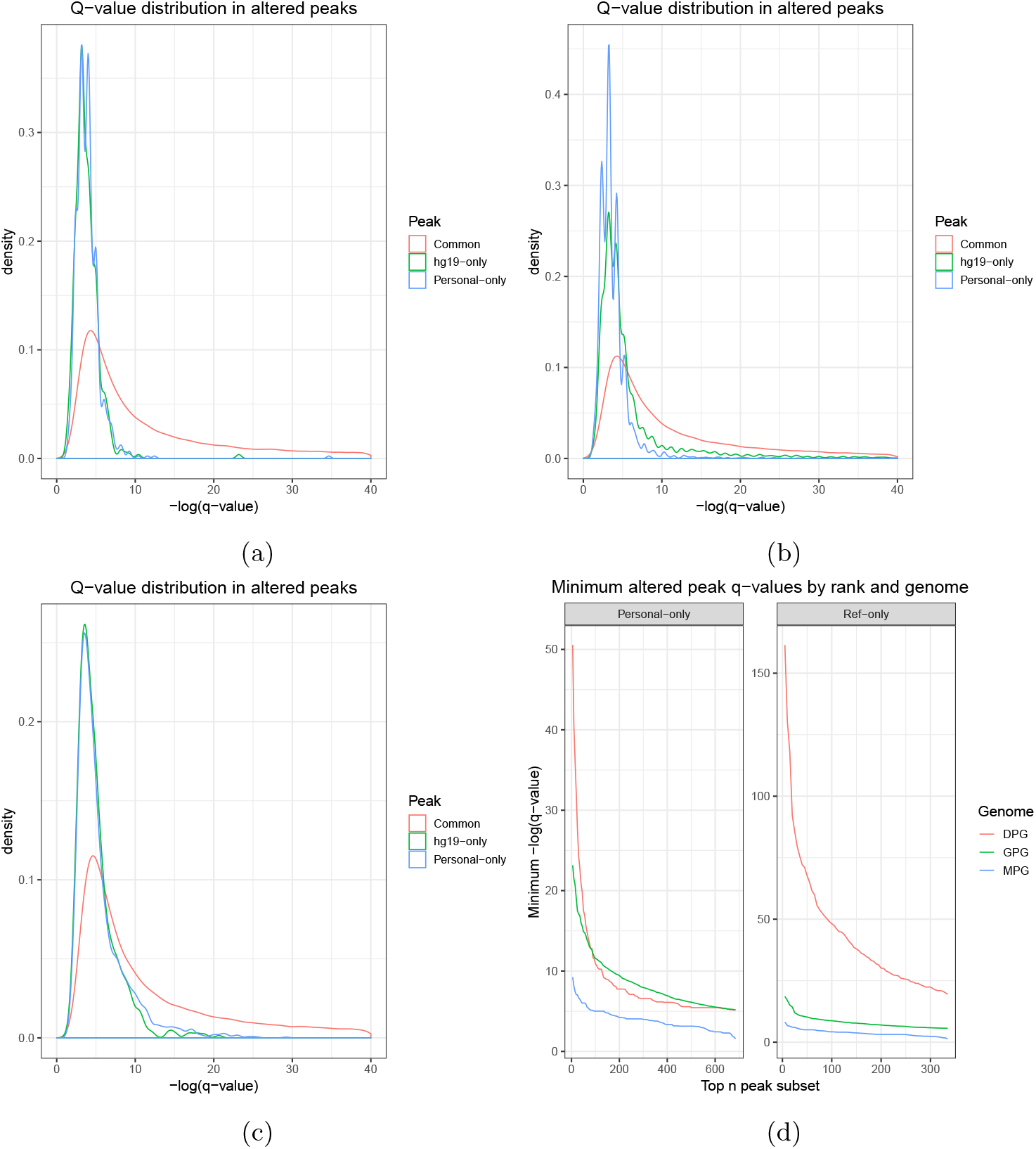
Q-value distributions for H3K27ac analogous to H3K4me1. a) In MPGs. b) In DPGs. c) In GPGs. d) Comparison by rank.

**Figure S4:**
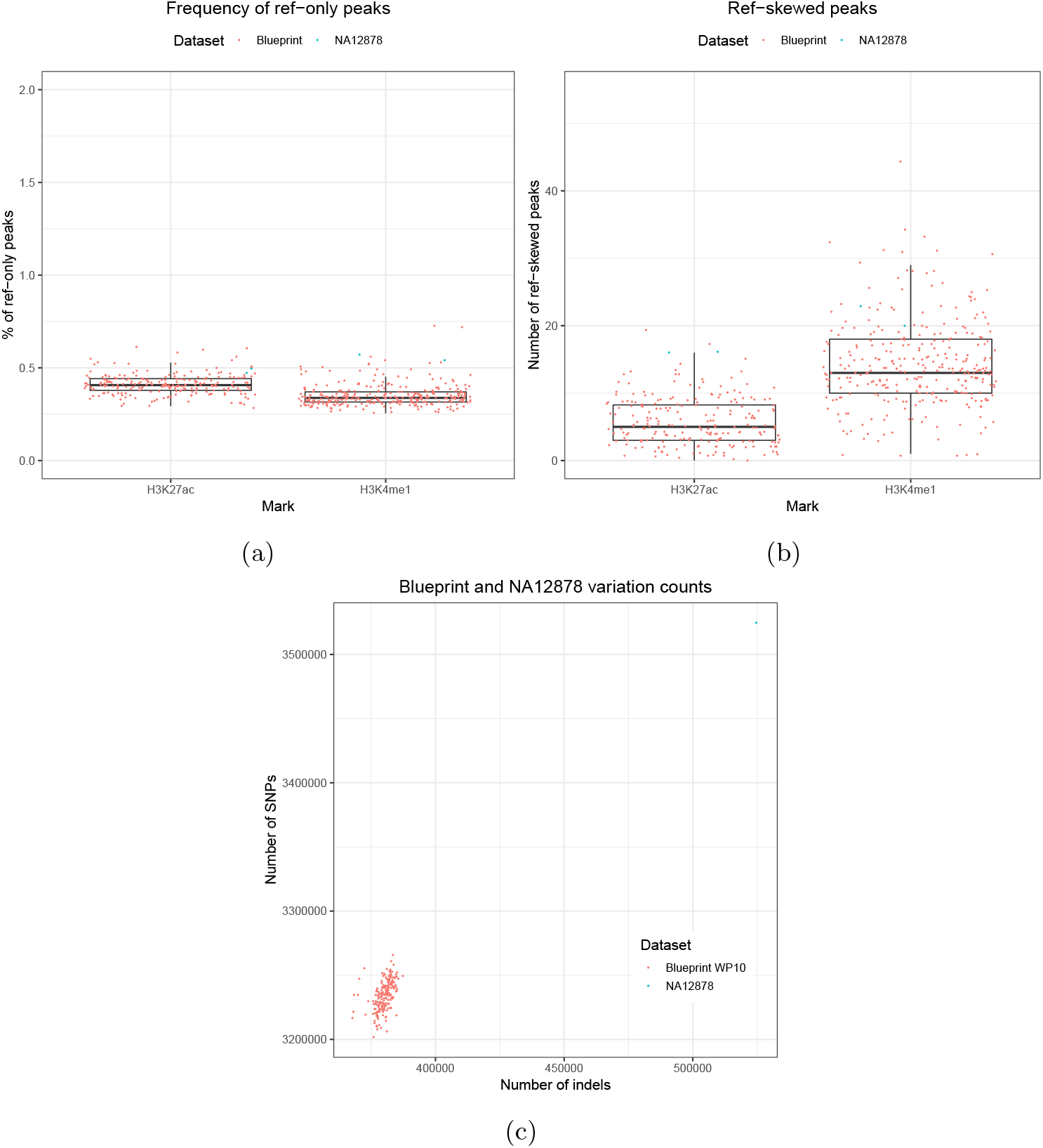
Comparison between NA12878 and Blueprint samples. a) Proportion of peaks that are called only in the reference. b) Number of peaks with higher coverage in the reference than the MPG. c) Difference in number of variation calls (SNPs and indels) between Blueprint samples and NA12878.

**Figure S5:**
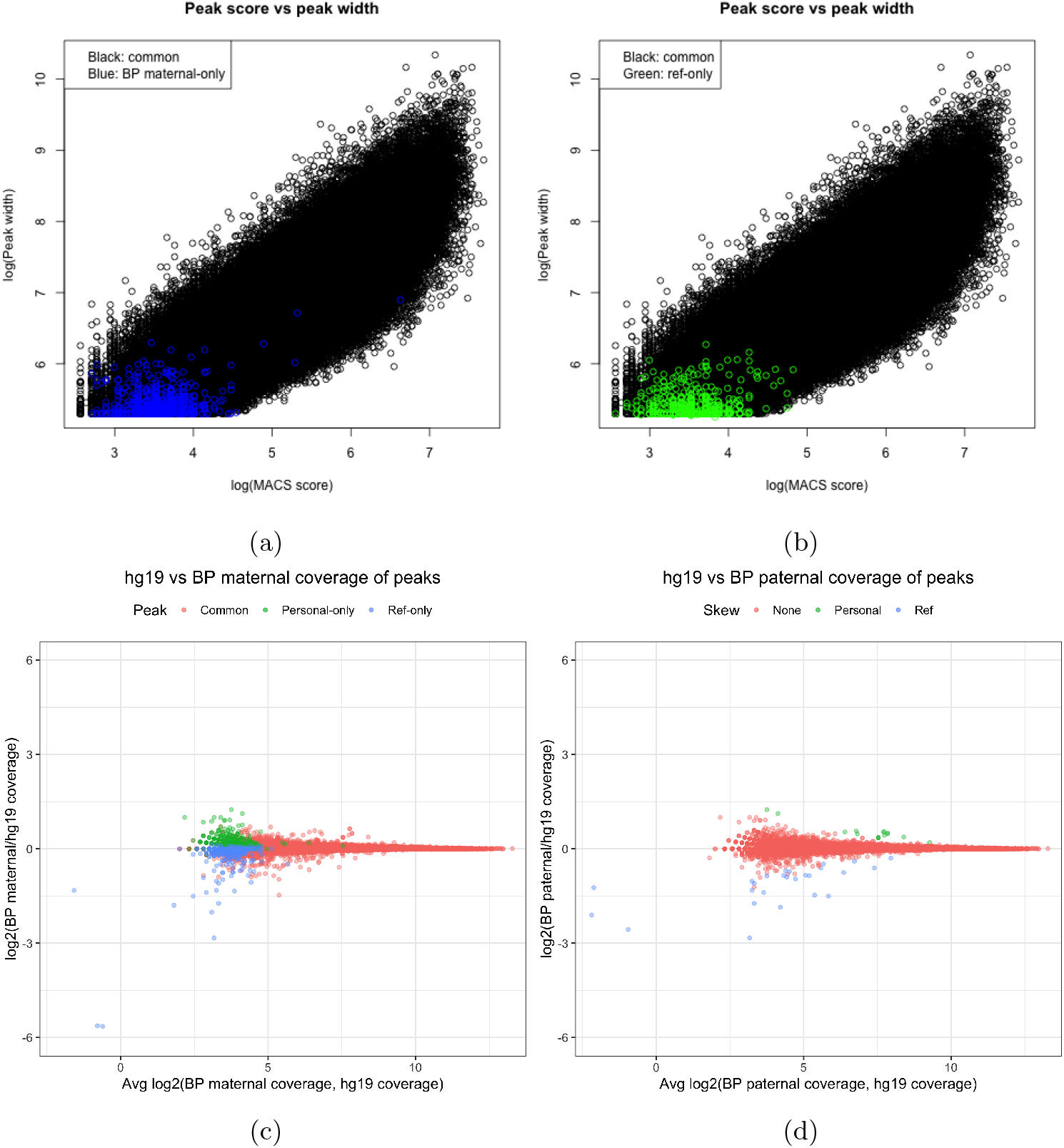
Characterization of AP calls in the maternal MPG of a typical Blueprint sample (H3K4me1).

**Figure S6:**
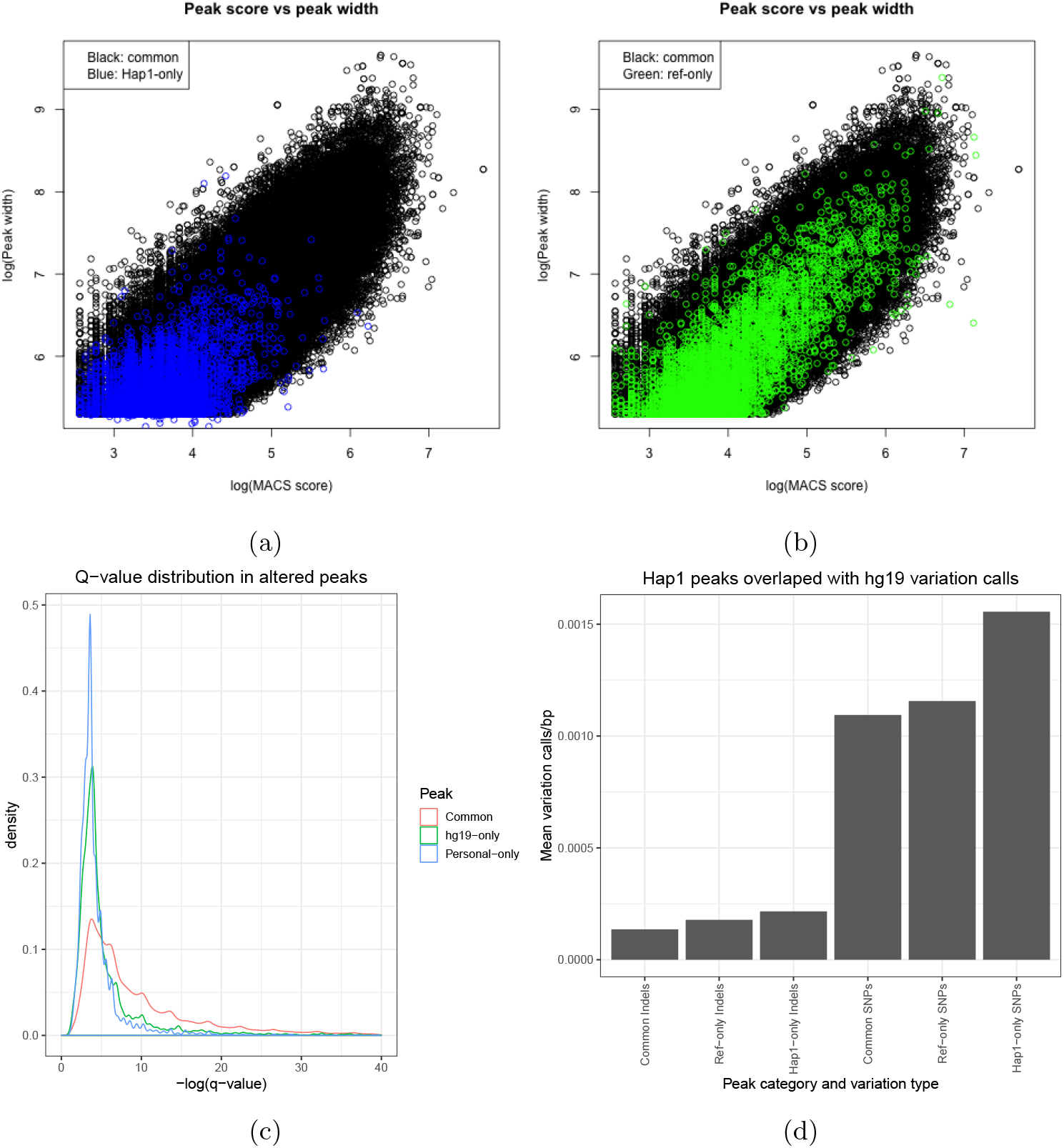
Confidence and width of the Hap1 DPG altered peak calls (H3K4me1). a) Peaks called only in Hap1 against the common peak background. b) Peaks called only in hg19 against the common peak background. c) Q-value distributions of the same peaks. d) Hap1-only peaks are enriched in hg19-relative variant calls relative to common peaks.

**Figure S7:**
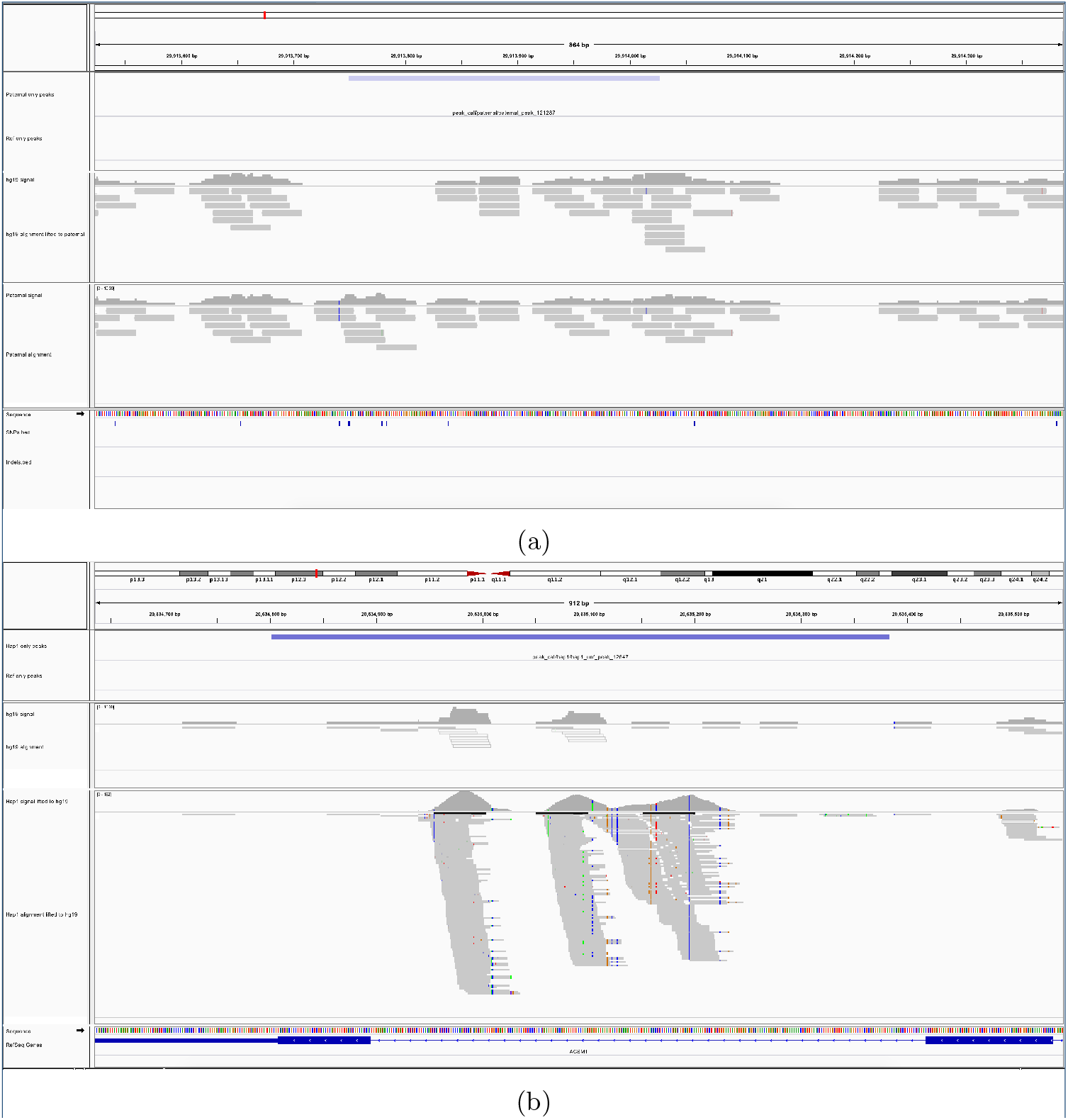
a) Typical personal-only peak in a NA12878 MPG. The small variations have a visible effect on only a few reads. (b) SD-free personal-only peak in the Hap1 DPG. Large scale changes in coverage become apparent with this approach.

**Figure S8:**
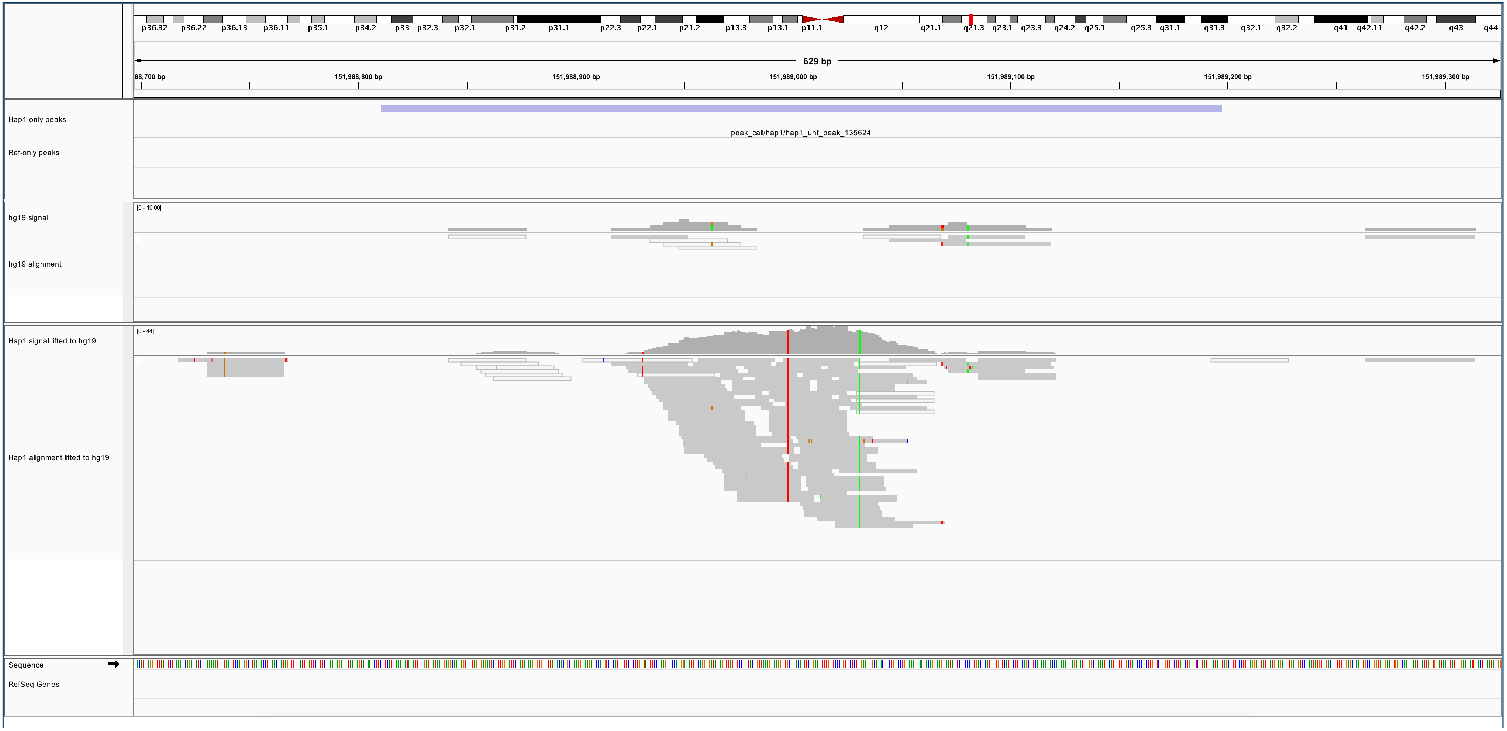
Hap1-only peak free of repeats and segmental duplications. Viewing this region in the UCSC genome browser shows overlaps with the hg19 self chain.

**Figure S9:**
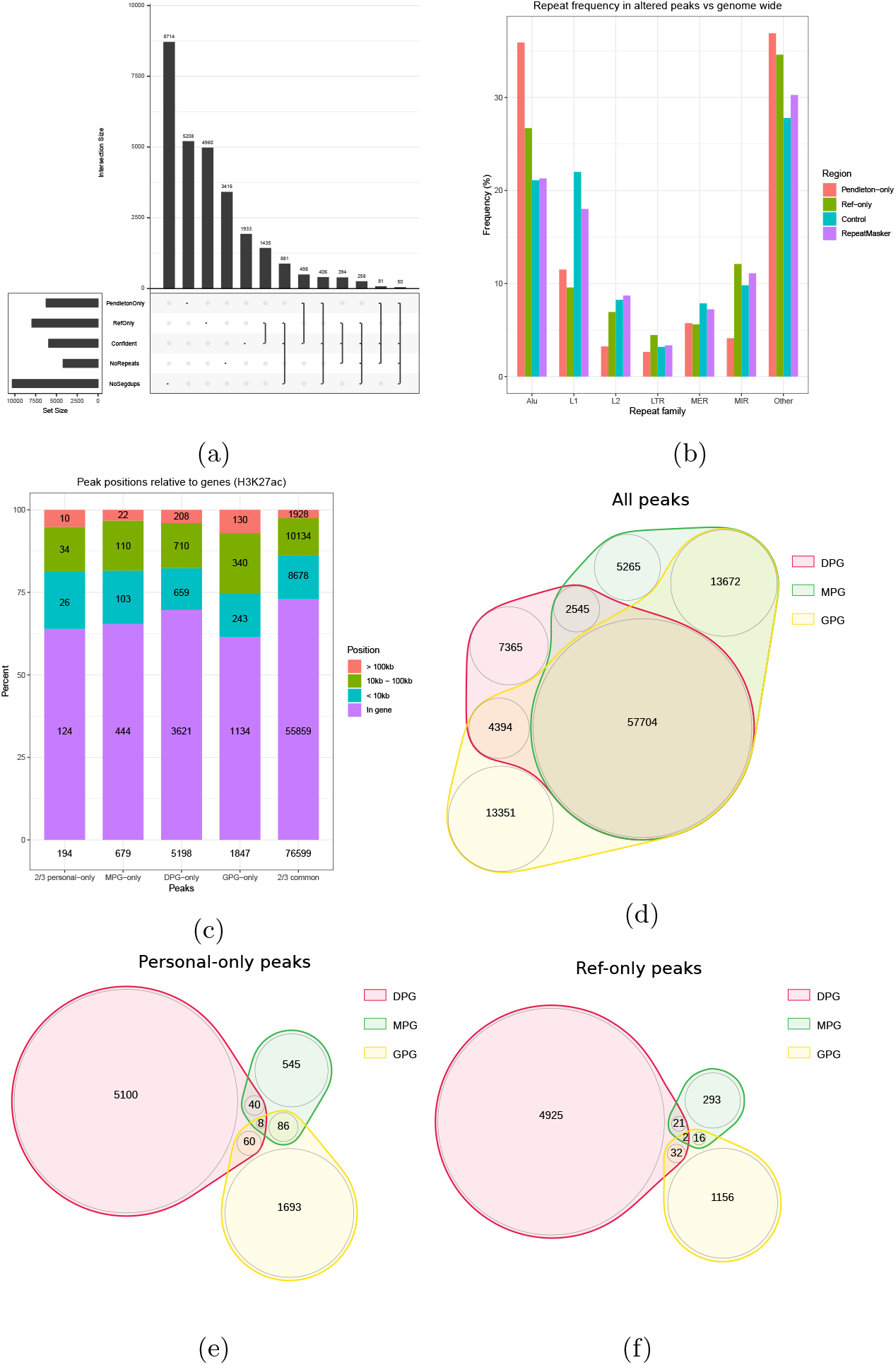
Plots for the H3K27ac mark that are analogous to H3K4me1. a) Summary of the overlap between altered peaks, confident peaks, repeats and segmental duplications. b) Frequency of repeat families within altered peaks compared to genome wide. The control is random intervals with the same width distribution as altered peaks. c) Distribution of gene relative positions of personal-only peaks among all genomes. The DPG is the Pendleton assembly. d) Overlap of all peak calls. e) Overlap of altered personal-only peak calls. f) Overlap of ref-only peak calls.

**Figure S10:**
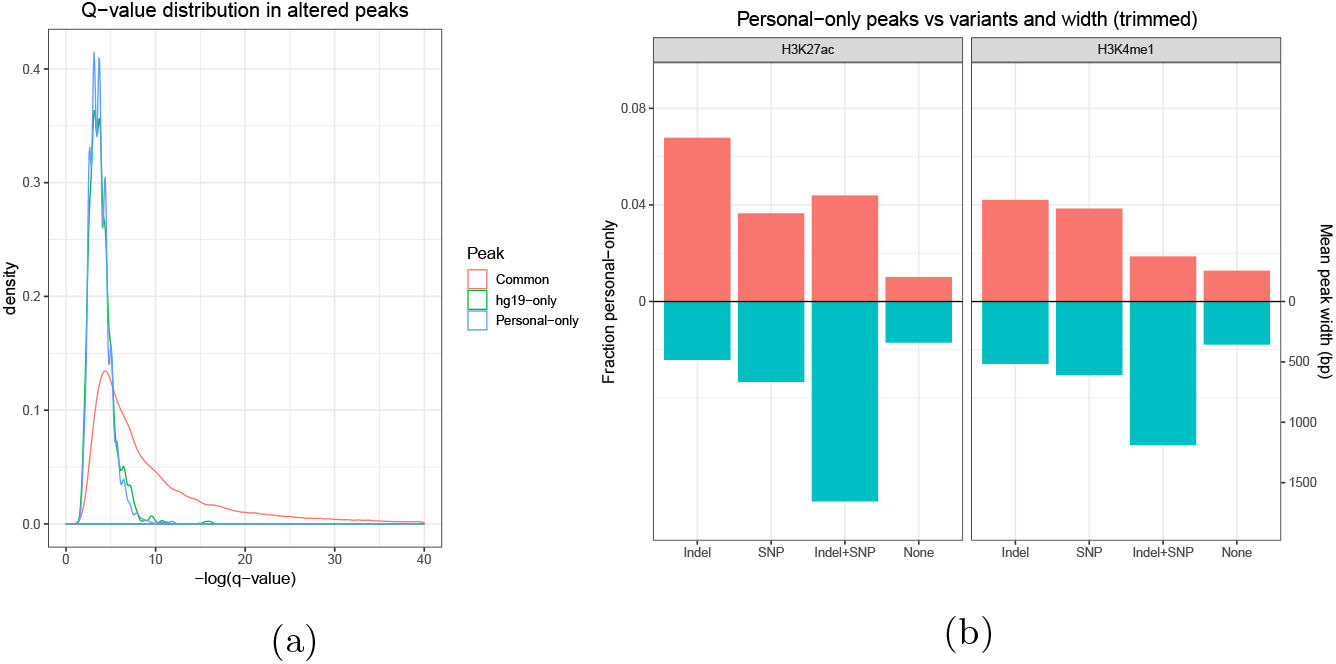
a) Q-value distribution of H3K4me1 altered peaks in GPGs. b) Replicated estimates of the probability that variants will cause an altered peak in GPGs.

**Figure S11:**
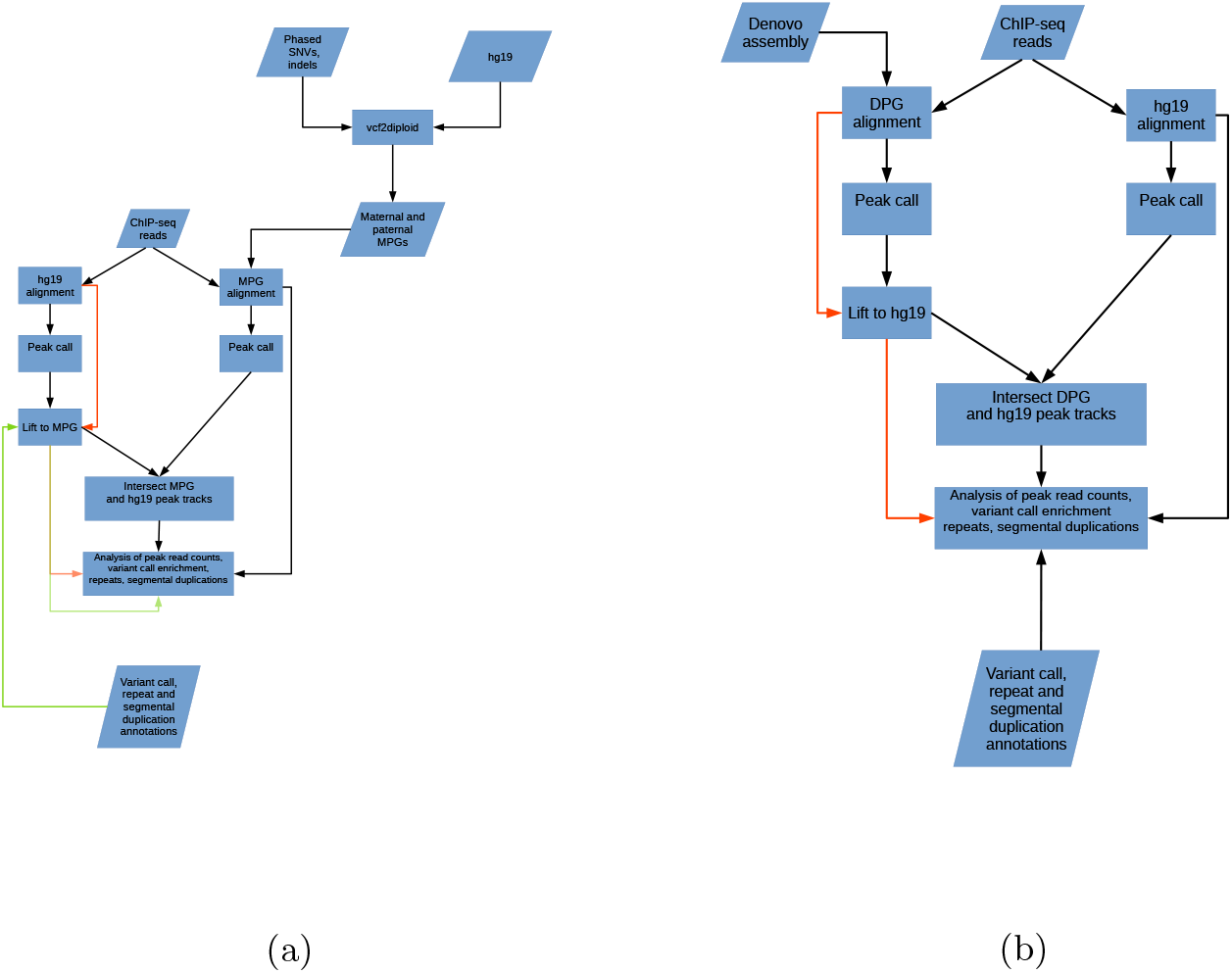
Flow chart of analysis for a) modified personal genomes and b) denovo personal genomes.

## Supporting tables

**Table S1:** a) Breakdown of WGS 36bp read alignment comparison between the reference and the NA12878 paternal MPG. The proportion of failed liftover, mapped/unmapped differences and unequal coordinate reads is 3.6%. This result is near identical in the both NA12878 MPGs. b) In the same comparison between the Hap1 DPG and the reference, the proportion is 9.79%. c) Comparing the alignments to the graph reference and the augmented NA12878 graph yields a proportion of 8.3% of reads with unequal mapping.

| Case (NA12878 MPG) | 36bp reads |  | Different |
| --- | --- | --- | --- |
| Failed liftover | 1947562 | 1.1% | Yes |
| Lifted coordinates | 167201030 | 90.4% | - |
| ↔ Equal in hg19, MPG | 162854003 | 88.0% | No |
| ↔ Unequal in hg19, MPG | 4347027 | 2.4% | Yes |
| Unmapped in hg19, mapped in MPG | 122163 | 0.07% | Yes |
| Mapped in hg19, unmapped in MPG | 46565 | 0.03% | Yes |
| Unmapped in hg19, unmapped in MPG | 15601588 | 8.4% | No |
| Total | 184918908 | 100% | 3.6% |

(a)
| Case (NA12878 DPG) | 36bp reads |  | Different |
| --- | --- | --- | --- |
| Failed liftover | 8968154 | 4.8% | Yes |
| Lifted coordinates | 160169017 | 86.6% | - |
| ↔ Equal in hg19, Hap1 | 152748687 | 82.6% | No |
| ↔ Unequal in hg19, Hap1 | 7420330 | 4.0% | Yes |
| Unmapped in hg19, mapped in Hap1 | 156895 | 0.08% | Yes |
| Mapped in hg19, unmapped in Hap1 | 1683155 | 0.91% | Yes |
| Unmapped in hg19, unmapped in Hap1 | 13941687 | 7.54% | No |
| Total | 184918908 | 100% | 9.79% |

(b)
| Case (NA12878 GPG) | 36bp reads |  |
| --- | --- | --- |
| Unequal mapping | 15378825 | 8.3% |
| Equal mapping | 169540083 | 91.7% |
| Total | 184918908 | 100 |

**Table S2:**
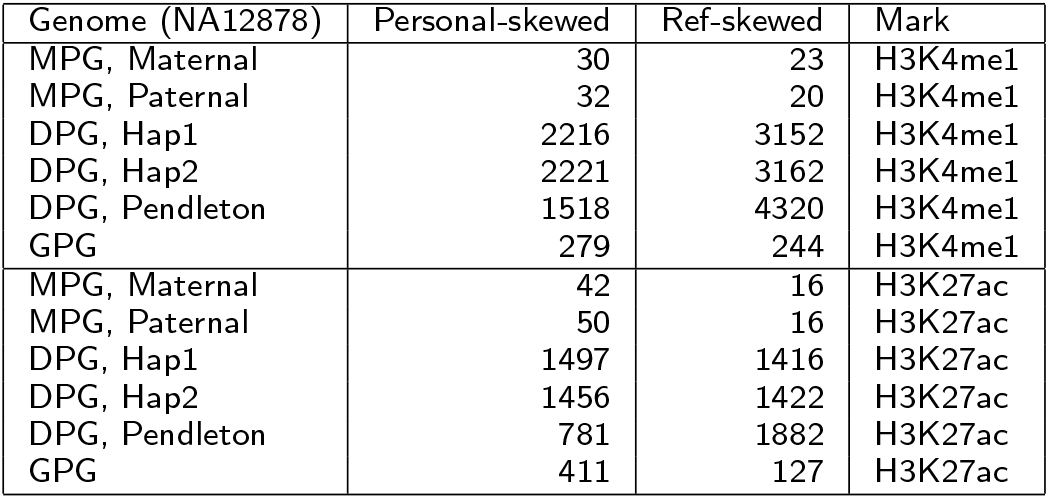
Number of peaks with skewed coverage in NA12878 MPGs and DPGs for H3K4me1 and H3K27ac marks. The read length across all rows was kept at 36bp.

| Genome (NA12878) | Personal-skewed | Ref-skewed | Mark |
| --- | --- | --- | --- |
| MPG, Maternal | 30 | 23 | H3K4me1 |
| MPG, Paternal | 32 | 20 | H3K4me1 |
| DPG, Hap1 | 2216 | 3152 | H3K4me1 |
| DPG, Hap2 | 2221 | 3162 | H3K4me1 |
| DPG, Pendleton | 1518 | 4320 | H3K4me1 |
| GPG | 279 | 244 | H3K4me1 |
| MPG, Maternal | 42 | 16 | H3K27ac |
| MPG, Paternal | 50 | 16 | H3K27ac |
| DPG, Hap1 | 1497 | 1416 | H3K27ac |
| DPG, Hap2 | 1456 | 1422 | H3K27ac |
| DPG, Pendleton | 781 | 1882 | H3K27ac |
| GPG | 411 | 127 | H3K27ac |

**Table S3:** Average number of altered peak calls in MPGs for the Blueprint samples for the same marks.

| Version | Mark | common | personal-only | ref-only |
| --- | --- | --- | --- | --- |
| MPG, Paternal | H3K4me1 | 129496 | 746 (0.6%) | 456 (0.4%) |
| MPG, Maternal | H3K4me1 | 129497 | 747 (0.6%) | 455 (0.4%) |
| MPG, Paternal | H3K27ac | 46830 | 336 (0.7%) | 196 (0.4%) |
| MPG, Maternal | H3K27ac | 46831 | 335 (0.7%) | 195 (0.4%) |

**Table S4:** a) Breakdown of WGS 100bp read alignment comparison between the reference and the NA12878 paternal MPG. The proportion of failed liftover, mapped/unmapped differences and unequal coordinate reads is 3.43%. This result is near identical in both NA12878 MPGs. b) In the same experiment with the Hap1 DPG and the reference, the proportion is 9.43%%. c) Comparing the alignments to the graph reference and the augmented NA12878 graph yields a proportion of 6.4% of reads with unequal mapping.

| Case (NA12878 MPG) | 100b reads |  | Different |
| --- | --- | --- | --- |
| Failed liftover | 2030979 | 1.1% | Yes |
| Lifted coordinates | 175123570 | 94.7% | - |
| ↔ Equal in hg19, MPG | 170876933 | 92.4% | No |
| ↔ Unequal in hg19, MPG | 4246637 | 2.3% | Yes |
| Unmapped in hg19, mapped in MPG | 39169 | 0.02% | Yes |
| Mapped in hg19, unmapped in MPG | 19307 | 0.01% | Yes |
| Unmapped in hg19, unmapped in MPG | 9632679 | 5.2% | No |
| Total | 184918908 | 100% | 3.43% |

(a)
| Case (NA12878 DPG) | 100bp reads |  | Different |
| --- | --- | --- | --- |
| Failed liftover | 9769845 | 5.3% | Yes |
| Lifted coordinates | 165311184 | 89.4% | - |
| ↔ Equal in hg19, Hap1 | 158400964 | 85.7% | No |
| ↔ Unequal in hg19, Hap1 | 6910220 | 3.7% | Yes |
| Unmapped in hg19, mapped in Hap1 | 49997 | 0.03% | Yes |
| Mapped in hg19, unmapped in Hap1 | 749507 | 0.4% | Yes |
| Unmapped in hg19, unmapped in Hap1 | 9038375 | 4.9% | No |
| Total | 184918908 | 100% | 9.43% |

(b)
| Case (NA12878 GPG) | 100bp reads |  |
| --- | --- | --- |
| Unequal mapping | 11776234 | 6.4% |
| Equal mapping | 173142674 | 93.6% |
| Total | 184918908 | 100 |

**Table S5:** Ref-only peaks with null coverage in the Hap1 DPG are extremely enriched in segmental duplications.

| Ref-only peaks | Number | Contain segmental duplications | Percent |
| --- | --- | --- | --- |
| Null coverage in Hap1 | 2547 | 1781 | 69.9 |
| ↔ Confident | 1490 | 1062 | 71.3 |
| Positive coverage in Hap1 | 4208 | 267 | 6.3 |
| ↔ Confident | 574 | 98 | 17.1 |

